# Proteins That Interact with the Mucin-Type Glycoprotein Msb2p Include Regulators of the Actin Cytoskeleton

**DOI:** 10.1101/786475

**Authors:** Aditi Prabhakar, Nadia Vadaie, Thomas Krzystek, Paul J. Cullen

## Abstract

Transmembrane mucin-type glycoproteins can regulate signal transduction pathways. In yeast, signaling mucins regulate mitogen-activated protein kinase (MAPK) pathways that induce cell differentiation to filamentous growth (fMAPK pathway) and the response to osmotic stress (HOG pathway). To explore regulatory aspects of signaling mucin function, protein microarrays were used to identify proteins that interact with the cytoplasmic domain of the mucin-like glycoprotein, Msb2p. Eighteen proteins were identified that comprised functional categories of metabolism, actin filament capping and depolymerization, aerobic and anaerobic growth, chromatin organization and bud growth, sporulation, ribosome biogenesis, protein modification by iron-sulfur clusters, RNA catabolism, and DNA replication and DNA repair. A subunit of actin capping protein, Cap2p, interacted with the cytoplasmic domain of Msb2p. Cells lacking Cap2p showed altered localization of Msb2p and increased shedding of Msb2p’s N-terminal glycosylated domain. Consistent with its role in regulating the actin cytoskeleton, Cap2p, and another Msb2p-interacting protein, Aip1p, were required for the enhanced cell polarization during filamentous growth. Our study identifies proteins that connect a signalling mucin to diverse cellular processes and may provide insight into new aspects of mucin function.

## INTRODUCTION

Mucins belong to a family of high-molecular weight glycoproteins. There are two types of mucins, transmembrane mucins and secreted mucins *^1-^*^5^. Transmembrane mucins are further divided into signaling and non-signaling mucins. The signaling mucins, which contain a cytosolic domain, regulate signal transduction pathways ^1, 4, 6^. In normal settings, mucins can typically line the oral and gastrointestinal cavity, where they engage in a variety of functions including protection and hydration of the host ^7^. When mis-regulated, mucins exhibit altered localization patterns and can contribute to unregulated signaling. Consequently, mucins are widely implicated in human diseases, such as inflammation and cancer ^1, 8–11^. Signaling mucins are structurally characterized by a heavily glycosylated extracellular N-terminal domain, a single pass transmembrane domain, and a cytoplasmic C-terminal domain ^12, 13^. The peptide backbone of the extracellular domain is rich in proline, threonine, and serine residues, which are typically found in tandem repeated sequences that are highly variable (PTS domain). The PTS domain distinguishes mucins from other large glycoproteins ^14–16^ and is heavily modified by multiple O-linked oligosaccharides ^17, 18^. This domain may be a convenient target for drug and vaccine development ^19^.

Transmembrane signaling mucins are found in a variety of organisms, including mammals, invertebrates, and microbes ^12, 20, 21^. In the budding yeast *Saccharomyces cerevisiae*, two well-known mucin-type glycoproteins, Msb2p and Hkr1p, have been identified ^20^. These proteins regulate MAPK pathways that respond to nutrients and osmotic stress. In response to nutrient limitation, yeast cells undergo a foraging response known as filamentous growth [or invasive/pseudohyphal growth ^22, 23^], which is typical of many fungal species ^24, 25^. Cells undergoing filamentous growth grow as branched filaments of elongated and connected cells ^26–29^. One of the pathways that regulates filamentous growth is a Cdc42p-dependent Mitogen Activated Protein Kinase (MAPK) pathway commonly referred to as the fMAPK pathway ^30, 31^. Msb2p functions at the plasma membrane to regulate the fMAPK pathway ^4, 5^. Msb2p is a single-pass transmembrane protein with a highly glycosylated N-terminal domain (1185 amino acids) connected to a cytoplasmic C-terminal signaling domain (97 amino acids) by a transmembrane domain. The cytoplasmic domain of Msb2p binds directly to Cdc42p ^20^. Cdc42p associates with the p21-activated kinase (PAK) Ste20p ^32, 33^ to regulate the fMAPK cascade [Ste11p, Ste7p, and Kss1p ^34^] that culminates in the phosphorylation/activation of transcription factors [Ste12p and Tec1p ^35^], which induce the expression of filamentation target genes ^36, 37^. The glycosylation of Msb2p is related to its signaling function ^38^. Under nutrient-limiting conditions, Msb2p is underglycosylated, which results in its proteolytic processing through a quality-control pathway called the Unfolded Protein Response (UPR) in the lumen of ER. The UPR up-regulates the expression of an aspartyl protease, Yps1p that processes Msb2p in its extracellular domain ^39, 40^. Proteolytic processing of Msb2p is required for activation of of the fMAPK pathway ^40^.

Another signaling mucin in yeast is Hrk1p, which regulates the Ste11p branch of the HOG pathway ^41^. The HOG pathway has two branches that converge on the MAPKK Pbs2p. Hrk1p does not regulate the fMAPK pathway ^42^. Moreover, overexpression of Msb2p and Hkr1p induce different sets of target genes ^42^. Thus, it is plausible that Msb2p preferentially regulates the fMAPK pathway, and Hkr1p preferentially regulates the HOG pathway. This may be an oversimplification, however, because Msb2p and Hrk1p can together regulate the HOG pathway ^41, 43^.

Despite the fact that much is known about the cleavage and activation mechanism of Msb2p (**Fig. 1**), relatively little is known about how the cytosolic domain connects to signal transduction pathways. It has previously been shown that Msb2p’s cytoplasmic domain binds to GTP-Cdc42p ^20^. More recently, Msb2p has been shown to interact with the Cdc42p adaptor, Bem1p ^44^. Whether these interactions represent the key interactions that tie Msb2p to Cdc42p-dependent pathways is not clear. Considering that mammalian mucins, like MUC1, interact with many different proteins to regulate cellular functions, one might also expect that Msb2p might have other biologically-relevant interactions that have not been explored. Here, we use protein microarray technology to identify new Msb2p interacting proteins. The proteins that associated with Msb2p have diverse cellular functions and may provide new connections between signaling mucins connect and different cellular processes.

**Figure 1.**
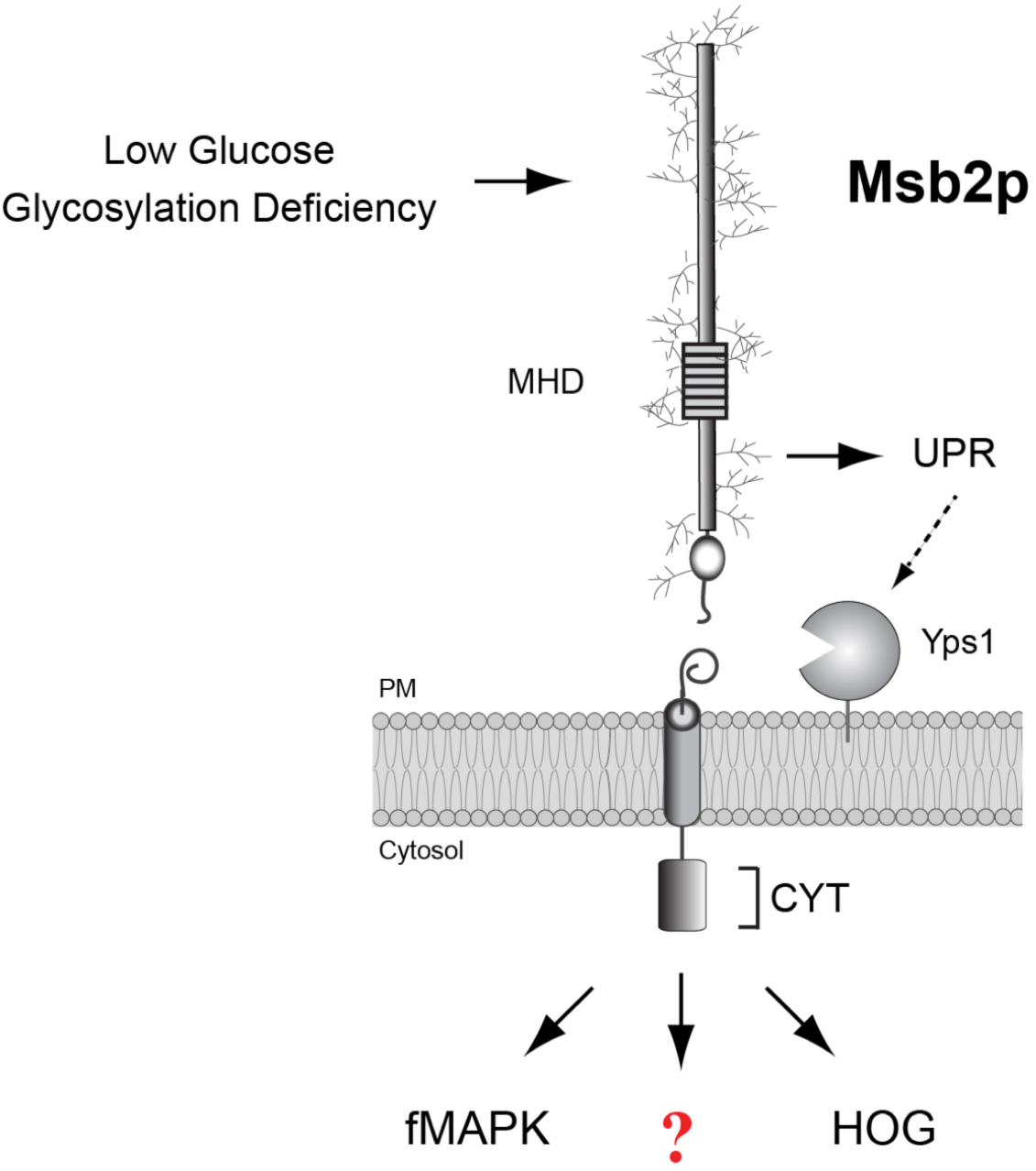
The mucin-like glycoprotein Msb2p. Msb2p shares a N-terminal mucin homology domain (MHD) with mammalian mucins. Msb2p is a single pass transmembrane protein and has a small cytoplasmic tail of 98 residues. Under-glycosylation of Msb2p in low nutrient environment triggers cleavage-dependent activation by the Unfolded Protein Response (UPR), which stimulates the filamentous growth MAPK pathway (fMAPK). Msb2p also regulates the High Osmolarity Glycerol response pathway (HOG). Red question mark, unknown interactions; PM, plasma membrane, CYT, cytosolic domain.

## MATERIALS AND METHODS

### Strains and Plasmids

Yeast strains are described in *Table 1*. Overexpression constructs were obtained from an ordered collection obtained from Open Biosystems ^45^. Gene disruptions and *GAL1* promoter fusions were made by PCR-based methods ^46, 47^ using plasmids provided by John Pringle (Stanford University, Palo Alto, CA). Some disruptions were created by the use of antibiotic resistance markers on cassettes *KanMX6* (PC1485) ^47^, *HYG* (PC2206) and *NAT* (PC2205) ^48^. Internal epitope fusions were created as described ^49^ using plasmids containing the 3XHA (PC1885) epitope. Strains containing TAP ^50^ and GFP ^51^ epitope fusions were provided by Michael Yu (University at Buffalo, Buffalo, NY). Gene disruptions were confirmed by PCR analysis and phenotype.

**Table 1.** Yeast strains used in this study.

| Name | Genotype | Reference |
| --- | --- | --- |
| PC471 | *MATa ura3-52 bud6::KIURA3<sup>a</sup>* | <sup>1</sup> |
| PC538 | *MATa ste4 FUS1-lacZ FUS1-HIS3 ura3-52* | <sup>2</sup> |
| PC539 | *MATa ste4 FUS1-lacZ FUS1-HIS3 ura3-52 ste12::URA3* | <sup>2</sup> |
| PC544 | *MATa ste4 FUS1-lacZ FUS1-HIS3 ura3-52 bni1::KIURA3* | <sup>1</sup> |
| PC551 | *MATa ste4 FUS1-lacZ FUS1-HIS3 ura3-52 pea2::URA3* | <sup>1</sup> |
| PC554 | *MATa ste4 FUS1-lacZ FUS1-HIS3 ura3-52 spa2::KIURA3* | <sup>1</sup> |
| PC948 | *MATa ste4 FUS1-lacZ FUS1-HIS3 ura3-52*<br>*msb2::KanMX6* | <sup>2</sup> |
| PC999 | *MATa ste4 FUS1-lacZ FUS1-HIS3 ura3-52 MSB2-*<br>*HA@500 aa* | <sup>2</sup> |
| PC1057 | *MATa ste4 FUS1-lacZ FUS1-HIS3 ura3-52 ste11::URA3* | <sup>1</sup> |
| PC1079 | *MATa ste4 FUS1-lacZ FUS1-HIS3 ura3-52 MSB2-HA*<br>*ste12::URA3* | <sup>3</sup> |
| PC1516 | *MATa ste4 FUS1-lacZ FUS1-HIS3 ura3-52 MSB2-HA<sup>Δ100-818</sup>* | <sup>2</sup> |
| PC3181 | *MATa ste4 FUS1-lacZ FUS1-HIS3 ura3-52 MSB2-HA*<br>*cap2::KIURA3* | This study |
| PC3184 | *MATa ste4 FUS1-lacZ FUS1-HIS3 ura3-52 MSB2-HA*<br>*gpp2::KIURA3* | This study |
| PC3190 | *MATa ste4 FUS1-lacZ FUS1-HIS3 ura3-52 MSB2-HA*<br>*mam33::KIURA3* | This study |
| PC3210 | *MATa ste4 FUS1-lacZ FUS1-HIS3 ura3-52 GAL-MSB2-*<br>*HA@500aa::KanMX6 cap2::KIURA3* | This study |
| PC3212 | *MATa ste4 FUS1-lacZ FUS1-HIS3 ura3-52 MSB2<sup>Δ100-818</sup>*<br>*cap2::KIURA3* | This study |
| PC3214 | *MATa ste4 FUS1-lacZ FUS1-HIS3 ura3-52 MSB2-HA*<br>*aip1::KIURA3* | This study |
| PC3216 | *MATa ste4 FUS1-lacZ FUS1-HIS3 ura3-52 MSB2-HA*<br>*nap1::KIURA3* | This study |
| PC3295 | *MATa ste4 FUS1-lacZ FUS1-HIS3 ura3-52 MSB2-*<br>*HA@500 aa CAP2-HA::KanMX6* | This study |
| PC3297 | *MATa ste4 FUS1-lacZ FUS1-HIS3 ura3-52 MSB2-HA*<br>*@500aa cap1::KIURA3* | This study |
| PC3299 | *MATa ste4 FUS1-lacZ FUS1-HIS3 ura3-52 MSB2-HA*<br>*cap2::NAT* | This study |
| PC3409 | *MATa ste4 FUS1-lacZ FUS1-HIS3 ura3-52 MSB2-HA*<br>*sac6::KIURA3* | This study |
| PC6928 | *MATa ste4 FUS1-lacZ FUS1-HIS3 ura3-52 nap1::NAT* | This study |
| PC6930 | *MATa ste4 FUS1-lacZ FUS1-HIS3 ura3-52*<br>*msb2::KanMX6 nap1::NAT* | This study |
| PC3128<sup>b</sup> | *MATa his3Δ1 leu2Δ0 met15Δ0 ura3Δ0 CAP2-TAP-*<br>*SpHIS3MX* | <sup>4</sup> |
| PC6743<sup>c</sup> | *MATa his3Δ1 leu2Δ0 met15Δ0 ura3Δ0 GLN1-GFP(S65T)-*<br>*SpHIS3MX<sup>d</sup>* | <sup>5</sup> |

All strains are Σ1278b background unless otherwise indicated.
- KI refers to *Kluyveromyces lactis*. - S288c background TAP collection from Ghaemmaghami S. et al 2003. - S288c background GFP collection from Huh W. K. et al 2003. - Sp refers to *Schizosaccharomyces pombe*

The pRS series of plasmids (pRS316) have been described ^52^. pET28b-*HIS*-*MSB2*^CYT^^20^ (PC1078), p*MSB2*-*GFP* ^40^ (PC1638 contains a C-terminal fusion of GFP to *MSB2*), p*GFP*-*MSB2* ^40^ (PC1638 contains an internal fusion of GFP to *MSB2*), YCp50-*STE11-4* ^53^ (PC1333), p*GAL*-*GFP*-*MSB2*@500 aa *CEN/URA* (*KanMX6*) (PC3406) ^40^ plasmids are published.

### Microbiological Techniques

Yeast and bacterial strains were manipulated by standard methods ^54, 55^. All experiments were carried out at 30°C unless otherwise indicated. ß-galactosidase assays were performed as described ^56^. Values represent the average of at least 3 independent trials. The mating-specific reporter *FUS1* was also used ^57^, which in cells lacking an intact mating pathway (*ste4*Δ) exhibits Msb2p- and fMAPK-dependent expression ^20^. *FUS1-HIS3* expression was used to confirm *FUS1-lacZ* reporter data and was measured by spotting equal amounts of cells onto synthetic medium lacking histidine and containing 4-amino-1,2,4-triazole. The single cell invasive growth assay ^58^ and plate-washing assay ^30^ were performed as described.

Budding pattern was based on established methodology ^59^ and was confirmed by visual inspection of connected cells. Mid-log cells were resuspended in water, and stained with Calcofluor white fluorescent brightener (Sigma-Aldrich Life Science and Biochemicals, St. Louis, MO) to a final concentration of 0.01% for 30 min, at 30°C. Cells were washed once in water. Bud scars were visualized by fluorescence microscopy.

### Microscopy

Differential-interference-contrast (DIC) and fluorescence microscopy of the GFP protein using FITC filter sets were performed using an Axioplan 2 fluorescent microscope (Zeiss) with a PLAN-APOCHROMAT 100X/1.4 (oil) objective (Zeiss). For most experiments, cells were visualized by resuspending in water at 25°C. Digital images were obtained with the Axiocam MRm camera (Zeiss). Axiovision 4.4 software (Zeiss) was used for image acquisition and analysis. Brightness and contrast of images was further adjusted in Adobe Photoshop.

### Immunoblot analysis

Detection of phosphorylated MAP kinase (P∼Kss1p) by immunoblot analysis was performed as described ^60–62^. Mid-log phase cells were harvested and pellets were washed once with water and stored at -80°C. Proteins were precipitated by trichloroacetic acid (TCA), separated by SDS-PAGE on 10% acrylamide gels and transferred to nitrocellulose membranes (AmershamTM ProtranTM Premium 0.45 µm NC, GE Healthcare Life sciences, 10600003). For blots to evaluate phosphorylated MAP kinase protein, membranes were incubated in 1X TBST (10 mM TRIS-HCl pH 8, 150 mM NaCl, 0.05% Tween 20) with 5% BSA. For other immunoblots, membranes were incubated in blocking buffer (5% nonfat dry milk, 10mM Tris-HCl pH8, 150mM NaCl and 0.05% Tween 20) for 16 h at 4°C. P∼Kss1p was detected using p44/42 antibodies (Cell Signaling Technology, Danvers, MA, 4370) at a 1:5,000 dilution. Kss1p was detected using α-Kss1p antibodies (Santa Cruz Biotechnology, Santa Cruz, CA; #6775) at a 1:5,000 dilution. α-HA antibody was used at a 1:5,000 dilution (Roche Diagnostics, 12CA5). α-GFP antibody was used at a 1:1,000 dilution (Roche Diagnostics, clones 7.1 and 13.1; Catalog# 11814460001). For secondary antibodies, goat anti-rabbit IgG-HRP antibodies were used at a 1:10,000 dilution (Jackson ImmunoResearch Laboratories, Inc., West Grove, PA, 111-035-144). Mouse α-Pgk1p antibodies were used at a 1:5,000 dilution as a control for total protein levels (Novex, 459250). Goat α-mouse secondary antibodies were used at a 1:5,000 dilution to detect primary antibodies (Bio-Rad Laboratories, Hercules, CA, 170-6516). Secondary incubations were carried out at 25°C for 1 h. ECL Plus immunoblotting reagent was used to detect secondary antibodies (Amersham Biosciences, Piscataway NJ). Msb2p secretion by colony immunoblot analysis has been described ^63^.

### Protein Microarray Analysis

Protein microarrays (Invitrogen Life Technologies; lot# 052404) containing 4,318 yeast proteins ^64^ were probed with the purified and biotinylated cytoplasmic domain of the Msb2p protein ^20^. The cytoplasmic domain (CYT) of the Msb2p protein was purified from *Escherichia coli* as a HIS-epitope fusion as described ^20^. 50µg of purified HIS-Msb2p^CYT^ was biotinylated using the yeast protoarray PPI kit (Invitrogen Life Technologies). Protein microarrays were probed following manufacturer’s instructions. Arrays were blocked with PBST buffer for 1h at 4°C with gentle shaking. The microarrays were incubated with biotinylated probes for 1.5h at 4°C without any shaking. Arrays were washed with probing buffer (5X PBS pH 7.4, 0.25% Triton X-100 25% glycerol) twice on ice and incubated with Streptavidin-Alexa Fluor 647 solution for 30 min on ice in the dark. The arrays were washed three times in the probing buffer at 4°C. The microarrays were centrifuged at 800 x g and dried at room temperature for 30 min. Dried arrays were scanned using GenePix 4000B (Molecular Devices Corporation).

The numerical data (GenePix result file) for the array was acquired using GenePix Pro 6.0 software (Molecular Devices Corporation) and processed using ProtoArray® Prospector v5.1 (Invitrogen Life Technologies). This software calculated statistical parameters including Z-factor (which measures the signal-to-noise ratio), Z-score (which indicates the number of standard deviations away from the mean signal value for all protein spots), replicate spot CV (coefficient of variation) and p-value for each spot. A protein was considered as a potential Msb2p^CYT^ interacting partner if it satisfied the following criteria: the protein signal in the experimental microarray after background subtraction was > 3-fold than the corresponding signal in the control microarray, the Z-factor was greater than 0.5, corresponding to > 2-fold increase in the signal relative to background, the Z-score was >3.0, the variation (CV) between replicate spots was less than 50%, and the p-value that measured the probability that the observed signal resulted from the distribution of negative control signals was <0.05. The candidates that met these criteria were visually confirmed as hits on the array. *Table S1* summarizes the raw and analyzed data for the protein microarray.

### Co-Immunoprecipitation Analysis Between Msb2p and Cap2p

Co-IPT analysis was based on previous methodology ^65^. Spheroplasts were generated as described ^40^. Cell pellets were washed in 1ml of 1.2 M sorbitol and centrifuged for 4 min at low speed as before. Spheroplasts were lysed in 1ml of ice-cold IPT buffer (50 mM Tris-HCl pH 8, 1mM EDTA, 50mM NaCl, 1.5 % Igepal CA-630) supplemented with 1x protease inhibitor cocktail EDTA free and 1mM phenylmethylsulfonyl fluoride (PMSF) before use. Lysates were centrifuged at 4°C for 20 min at 15,800g. Supernatants were precleared by incubating with 30 ul of Immunopure immobilized Protein G plus on end-to-end rotator for 30 min. Beads were pelleted at 4°C for 2 min at 2,500 X g. Protein concentration of pre-cleared supernatants was determined by the Bradford Coomassie Blue assay. Five to seven mg of protein were IPed with poly-clonal antibody in 1:100 dilution for 1 h, and complexes were pulled down by adding 30 ul of protein G and incubating for 1 h. IPT complexes were pelleted at 2,500 X g for 1 min. Beads were washed in IPT buffer 3 times for 1 min. 50 ul of lysis buffer (8 M Urea, 5 % SDS, 40 mM Tris-HCl pH 6.8, 0.1 M EDTA, 0.4 mg/ml Bromophenol blue and 1 % ß-mercaptoethanol) was added to beads, and IPT complexes were released from beads by boiling for 10 min. Beads were removed by centrifuging for 1 min at 2,500 X g after which the pellet was discarded. Supernatants (protein complexes) were immediately processed by SDS-PAGE or frozen at -80°C.

### Pull downs between HIS-Msb2p^CYT^ and Cap2p-TAP

*E. coli* BL21λDE3 cells expressing pET28b-HIS-Msb2p^CYT^ (PC6071) were grown to mid-log phase and induced with 1mM of IPTG for 3h. Cell pellets were resuspended in lysis buffer [PBS supplemented with 1x protease inhibitor cocktail EDTA free and 1mM PMSF] and sonicated. Cell lysates were centrifuged at 14000 x g for 30 min, and cleared supernatants were incubated with 3 ml of Talon resin slurry (Clontech, Mountain View, CA) for 1 h. Cell homogenates were loaded onto Poly-Prep chromatography columns (Bio-Rad, Hercules, CA). The beads were washed three times, and the immobilized HIS-Msb2p^CYT^ beads were incubated with Cap2p-TAP cell lysates for 1 h. The homogenates were loaded onto columns, and protein complexes were eluted in PBS buffer containing 200mM imidazole (Sigma-Aldrich, St. Louis, MO 63103).

### Pull downs between HIS-Msb2p^CYT^ and Gln1p-GFP

Yeast cells harboring Gln1p-GFP (PC6743) from the GFP collection ^51^, or control cells (PC6021) were grown to mid-log phase in 800 ml YEPD medium at 30°C. Cells were harvested by centrifugation, at 4,000 rpm for 15 min at 4°C. The cell pellet was stored at -80°C.

400 ml of *E. coli* BL21λDE3 cells expressing pET28b-HIS-Msb2p^CYT^ (PC6071) were induced in 2XYT medium at an O.D._600_ = 0.6 with 0.3 mM IPTG at 37°C for 4 h. Cells were harvested by centrifugation at 4,000 rpm for 15 min at 4°C. Cells pellets were stored at -80°C. As a control, 400 ml of control *E. coli* BL21λDE3 cells lacking the plasmid pET28b-HIS-Msb2p^CYT^ were harvested at log phase.

Cells were resuspended in 40 ml lysis buffer [TRIS pH8, 100mM NaCl, 0.01% βME, 1mg/ml lysozyme, 1mM PMSF, 0.1% Triton X-100] and sonicated 30 sec on, 30 sec off for 15 min on ice. A portion of the cell extract was removed (*E. coli* WCE) and the remaining extract was centrifuged at 15,000 rpm for 15 min at 4°C. The supernatant was collected and mixed with 3.5 ml of Talon resin slurry (Clontech, Mountain View, CA), which had been pre-equilibrated in lysis buffer [TRIS pH8, 100mM NaCl, 0.01% βME]. Extracts and beads were incubated for 1 h at 4°C with end-over-end rotation. Extracts containing beads were added to a 25 ml Poly-Prep chromatography columns (Bio-Rad, Hercules, CA). Column was washed with 10 ml lysis buffer twice. Supernatants from yeast extracts were immediately added to the column. Yeast supernatants were made by resuspending cell pellets in 15 ml lysis buffer, and 7 ml of glass beads were added. Cells were vortexed at 4°C for 5 X 1 min pulses, with 1 min rest between cycles. A portion of the extract was set aside (yeast WCE). Cell debris was removed by centrifugation at 15,000 rpm for 15 min at 4°C. The supernatants were incubated with Msb2p^CYT^ or control *E. coli* extract coated beads for 30 min at 4°C with end-over-end rotation. Extracts containing beads were added back to the column and allowed to flow through the column with a gravity flow at an approximate rate of 0.5 ml/min. The column was then washed twice with 10 ml lysis buffer. Fractions were eluted from the column in elution buffer containing imidazole (10 mM, 20 mM, 100 mM, and 500 mM). WCEs, flow-through and wash fractions were collected and examined by SDS-PAGE analysis.

## RESULTS

### Identification of Msb2p Interacting Proteins by Protein Microarray Technology

Protein microarray technology (e.g. proteome chips) is an established technology to explore protein function ^64, 66–68^. Proteins microarrays can identify new protein binding partners ^69^. Protein chips can also identify post-translational modifications ^70, 71^ and interactions between proteins and DNA ^72^, RNA ^73^ and lipids ^74^. We used protein microarrays to identify proteins that interact with the cytoplasmic domain of Msb2p (**Fig. 1**, Msb2p^CYT^). To identify protein interactions, an epitope-tagged version of the cytoplasmic domain of Msb2p (HIS-Msb2p^CYT^) was expressed in *Escherichia coli* and purified by affinity chromatography. Approximately 50 µg of purified HIS-Msb2p^CYT^ protein was biotinylated *in vitro* and incubated with yeast ProtoArray protein microarrays, which contain 4,318 immobilized yeast proteins spotted in duplicate on a slide. Interactions were detected using Alexa Fluor® 647-streptavidin, which binds to biotinylated Msb2p. A control microarray was probed with an *E. coli* extract that was also biotinylated but came from cells that did not contain the pET28b-HIS-Msb2p^CYT^ plasmid.

The protein microarray probed with the HIS-Msb2p^CYT^ showed a number of potential interactions (**Fig. 2A**, *Table S1*). The protein microarray data was examined by additional bioinformatics analysis. The signal intensity of each spot was expressed as the fold increase between the experimental and control microarrays after background subtraction (*Table2*, Corrected Signal). The cut-off for p-values was set to < 0.05 (*Table 2*, p-value). The normalized score or Z-score was calculated as the number of standard deviations of a signal compared to the mean signal of all spots on the array. A Z-score of >3 was used, which represents a 99% cut off (**Fig. 2A**, red bar; *Table 2*, Z-score). Eighteen proteins qualified as Msb2p^CYT^-interacting proteins based on this criteria (*Table 2*).

**Figure 2.**
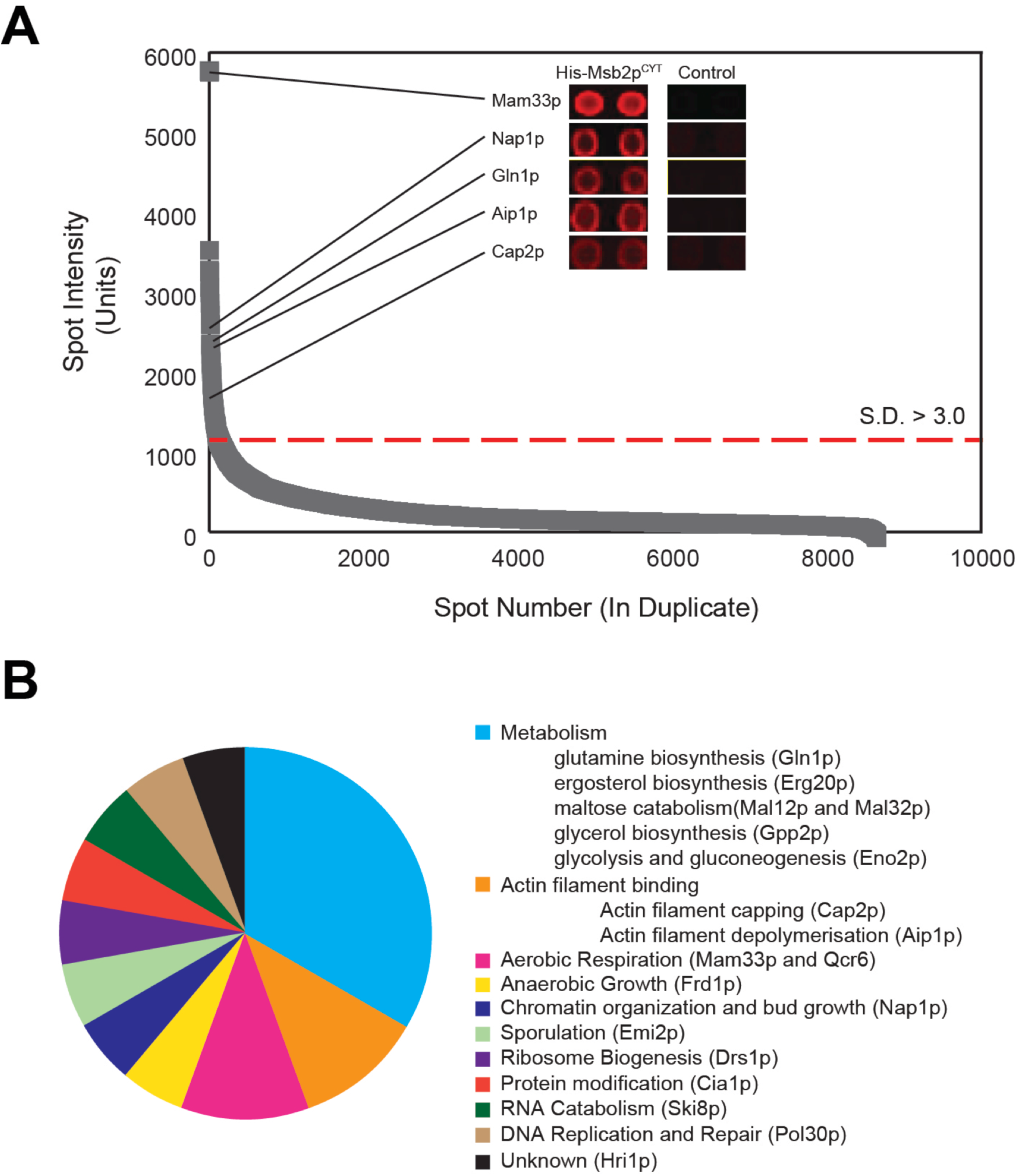
Msb2p-interacting proteins identified by the protein microarray. A) A protein microarray containing ∼4,000 yeast proteins spotted in duplicate was probed with a purified epitope-tagged version of the C-terminus of Msb2p (HIS-Msb2p^CYT^) that was biotinylated *in vitro*. The curve shows protein interactions ranked by signal intensity. The signal intensity of a spot was calculated by subtracting the background intensity for each spot. Potential ‘hits’ were identified using ProtoArray® Prospector v5.1 (Invitrogen Life Technologies). Inset, examples of spots that showed interaction with HIS-Msb2p^CYT^ in the experimental microarray (right) compared to the corresponding region in the control microarray (left). The dashed red line represents the cutoff for statistically significant interactions (>3.0 S.D.). See *Table S1* for the raw data. (**B**) Pie chart of Msb2p interacting proteins. The proteins are separated by GO terms obtained from *Saccharomyces* Genome Database.

**Table 2.**
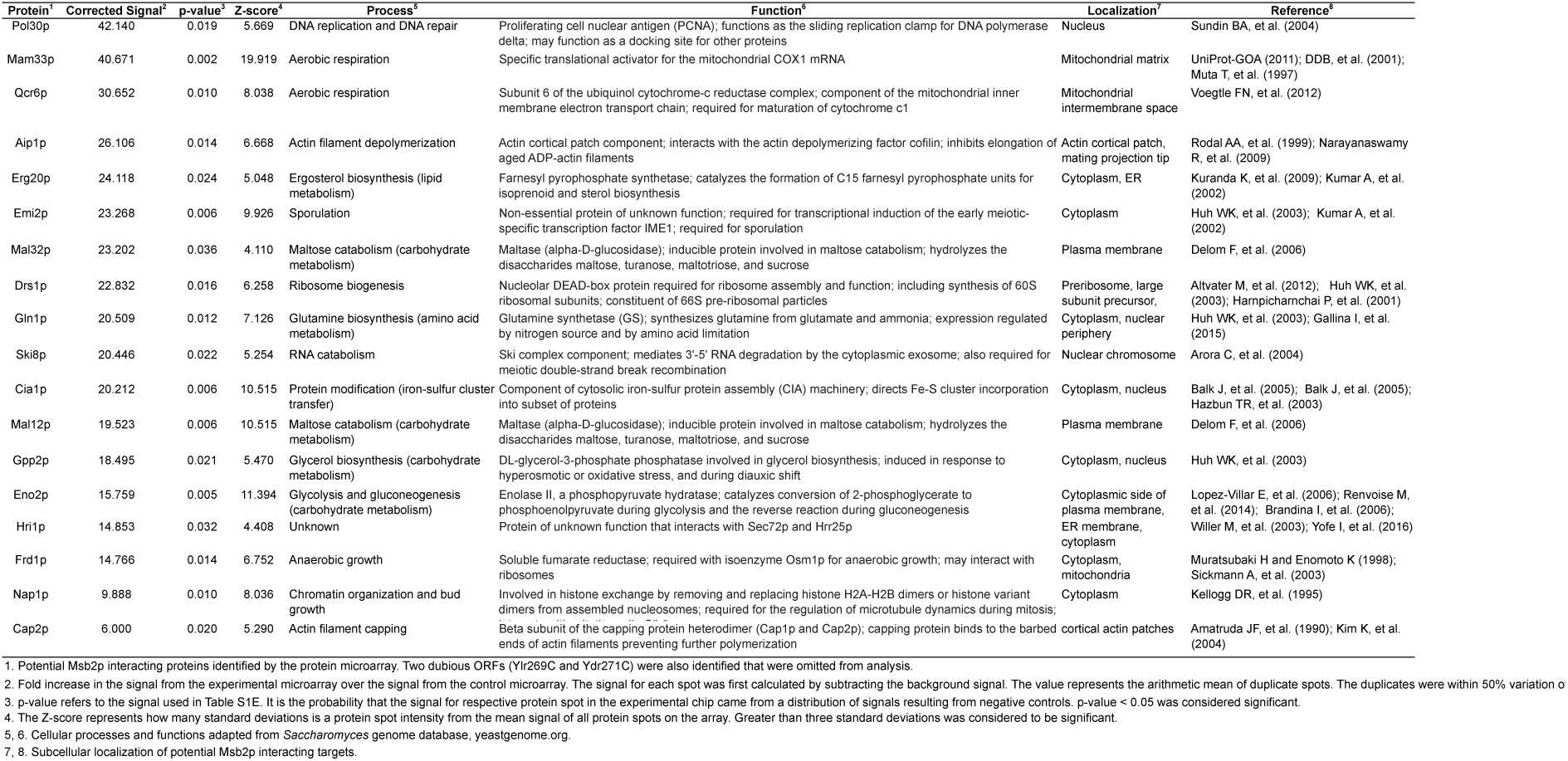
List of potential Msb2p interacting proteins identified by protein microarray.

### Functional Characterization of Msb2p-Interacting Proteins

Msb2p interacts with proteins that regulate the fMAPK pathway, including Sho1p ^20, 40^, Cdc42p ^20^, Opy2p ^75^, Mig1p ^76^, Mig2p ^76^ and Bem1p ^44^. None of these interactions were detected by the microarray. Msb2p and Sho1p interact by their transmembrane regions ^20, 40^ and would not be expected to be identified by this approach. Msb2p interacts with the GTP-bound conformation of Cdc42p ^20^. This conformation would not be favored on the microarray in the absence of GTP. Opy2p was not spotted on the microarray. Although present on the microarray, Mig1p, Mig2p, and Bem1p were not identified. The protein microarray may not capture all functionally relevant Msb2p interactions; thus, we focused on the interactions detected by this method.

Functional analysis by GO term analysis ^77, 78^ showed that the Msb2p-interacting proteins comprised functional groups including metabolism (Gln1p, Erg20p, Mal12p, Mal32p, Gpp2p and Eno2p), actin filament capping and depolymerization (Cap2p and Aip1p), aerobic (Mam33p and Qcr6p) and anaerobic growth (Frd1p), chromatin organization and bud growth (Nap1p), sporulation (Emi2p), ribosome biogenesis (Drs1p), protein modification by iron-sulfur clusters (Cia1p), RNA catabolism (Ski8p), and DNA replication and DNA repair (Pol30p) (**Fig. 2B***, Table 2*). Most of the proteins have been shown to localize to the cytoplasm or plasma membrane where Msb2p is known to reside (*Table 2*, Localization).

To determine whether newly identified Msb2p-interacting proteins regulate filamentous growth, which is regulated by Msb2p, isogenic strains carrying non-essential deletion mutations were constructed in the Σ1278b background (for *mam33*Δ, *aip1*Δ, *gpp2*Δ and *nap1*Δ; *Table 3*, Gene Deletion Mutant) or acquired from the nonessential Σ1278b deletion collection ^79^ (for *qcr6*Δ, *emi2*Δ, *mal32*Δ, *mal12*Δ, *frd1*Δ). The effect of protein over-expression on filamentous growth was also examined for some candidates (Pol30p, Erg20p, Ski8p, Mam33p, Hri1p, Aip1p, Eno2p, Frd1p and Nap1p; *Table 3*, Overexpression) using a genome-wide overexpression collection ^45^. The overexpression collection allowed evaluation of the role of several essential proteins in regulating filamentous growth (including Pol30p, Erg20p, Ski8p). Six Msb2p-interacting proteins impacted filamentous growth when deleted or overexpressed: Cap2p, Aip1p, Nap1p, Mam33p, Mal32p and Gln1p (*Table 3*). In particular, Cap2p, Aip1p, Mal32p and Gln1p were positive regulators of filamentous growth, while Nap1p and Mam33p were negative regulators of filamentous growth (*Table 3*; IG, Invasive Growth). The majority of these proteins comprised two functional categories (**Fig 2B**; *Table 2*; metabolism and actin filament capping and depolymerization) and were examined in more detail. Other Msb2p interacting proteins may impact Msb2p function in the HOG or other pathways ^41, 43, 44, 80, 81^ but were not explored here.

**Table 3.**
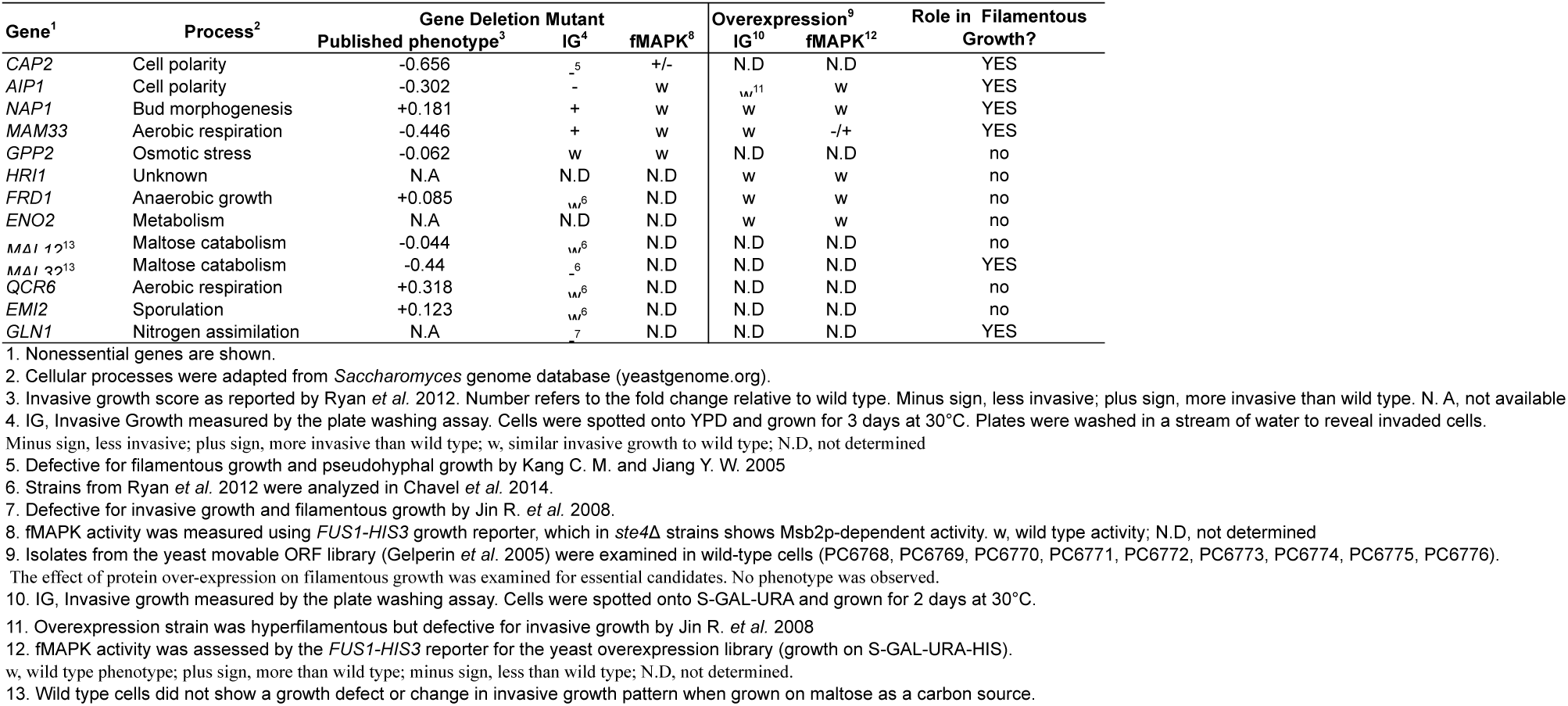
Functional analysis of Msb2p-interacting proteins identified by protein microarray.

### Msb2p and Cap2p interact in vitro and in vivo

Given that cytoskeleton-organizing proteins (Cap2p and Aip1p) were identified as potential Msb2p-interacting proteins, we became interested in exploring how the actin cytoskeleton might impact mucin function. Of the two actin-filament binding proteins, Cap2p showed the stronger phenotype in the secondary analysis compared to Aip1p and was explored in more detail. Cap2p is the ß-subunit of actin-capping protein that forms a heterodimer with Cap1p. Actin-capping protein binds to the barbed end of actin filaments to prevent their elongation ^82–86^. We first tested whether Msb2p and Cap2p interact by *in vitro* pull down analysis. Cap2p-TAP, expressed in yeast cells from its endogenous promoter ^50^, was tested for association with beads coated with HIS-Msb2p^CYT,^ expressed and purified from *E. coli*. Cap2p-TAP associated with HIS-Msb2p^CYT^ by *in vitro* pull downs (**Fig. 3A**). This result validates the protein microarray data identifying Cap2p and demonstrates that Msb2p and Cap2p interact *in vitro*.

**Figure 3.**
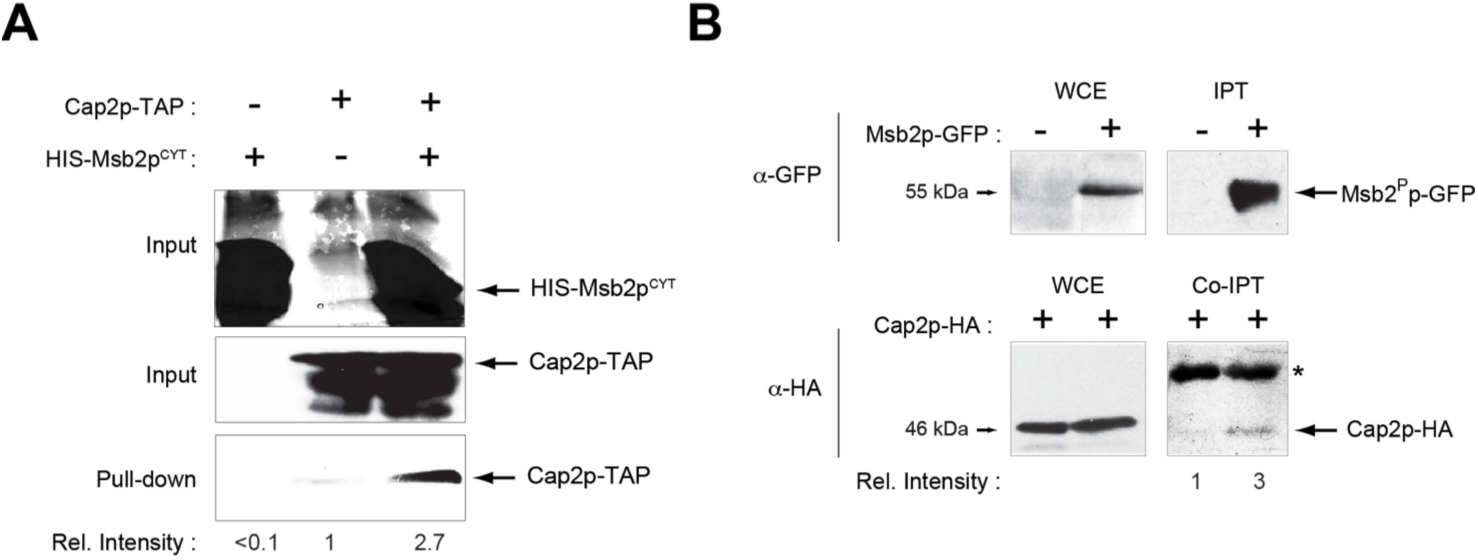
Actin-binding protein Cap2p interacts with Msb2p. (**A**) HIS-Msb2p^CYT^ and Cap2p-TAP associate by pull down analysis. (**B**) Msb2p-GFP and Cap2p-HA associate by co-IPT analysis. For panels A and B relative intensity is the fold increase in band intensity determined by ImageJ analysis.

To determine whether the interaction between Msb2p and Cap2p also occur *in vivo*, co-immunoprecipitation (co-IPT) analysis was performed. Full-length Msb2p is processed by proteolytic processing ^40^. A Msb2p-GFP fusion expressed from its endogenous promoter migrated at the size of the processed form of the protein (**Fig. 3B**, 56 kDa, Msb2^P^p-GFP). Msb2^P^p-GFP precipitated with anti-GFP antibodies was abundantly enriched by IPT analysis. IPT of Msb2^P^p-GFP co-immunoprecipitated Cap2p-HA (**Fig. 3B**). Thus, as expected from the above results, Cap2p interacts with a portion of the Msb2p protein that includes the cytosolic domain. Both the *in vitro* and co-IPT analysis showed that a sub-stoichiometric amount of Cap2p was precipitated by Msb2p. Collectively, the results show that Msb2p interacts with Cap2p by *in vitro* and *in vivo* tests.

### Role of Cap2p in Regulating Msb2p Trafficking and Filamentous Growth

Cap2p and Aip1p are components of actin cortical patches. Actin patches are important for bud growth and endocytosis, and Cap2p has a specific function in the regulation of endocytosis ^87, 88^. Msb2p is a single pass transmembrane protein that is delivered to the plasma membrane and turned over in the lysosome (vacuole in yeast) through the secretory pathway ^39, 75^. Cap2p might influence the trafficking of Msb2p in the secretory pathway. A functional GFP fusion in the N-terminus of Msb2p (driven by a galactose-inducible promoter) was used to evaluate the localization of Msb2p in wild-type cells and the *cap2*Δ mutant. In wild-type cells, GFP-Msb2p was localized to the cell cortex and the lumen of the vacuole [**Fig. 4A**, wild type ^40^]. Although it has not been previously reported, the highly expressed GFP-Msb2p was also enriched at the mother-bud neck. In the *cap2*Δ mutant, GFP-Msb2p was mislocalized in some cells (∼27%), where the protein had a reticulated pattern (**Fig. 4A**, *cap2*Δ). The reticulated pattern of GFP-Msb2p in the *cap2*Δ mutant was higher than seen in wild-type cells (**Fig. 4B**).

**Figure 4.**
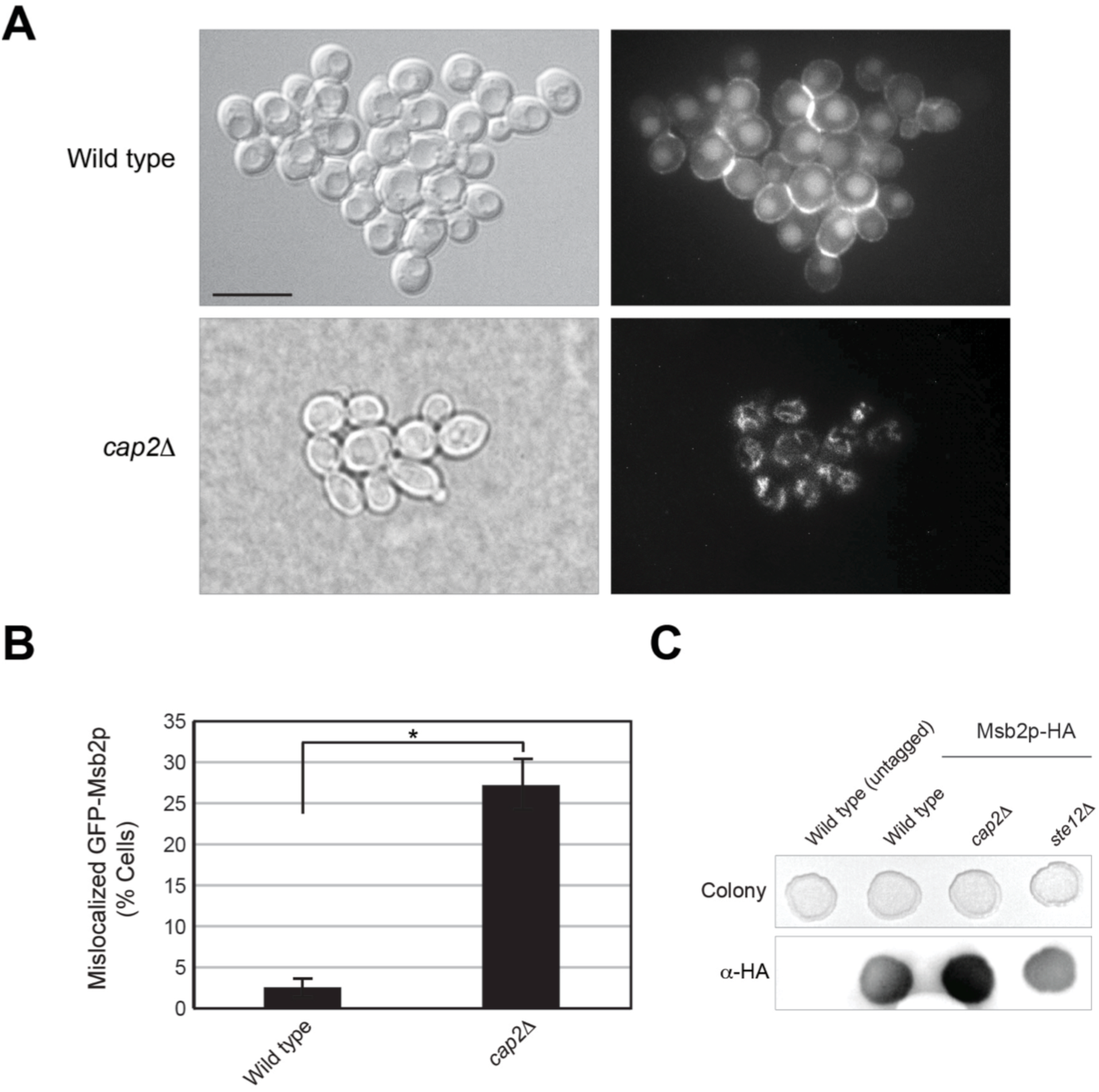
**Role of Cap2p in Msb2p trafficking and shedding**. (**A**) Micrographs of wild-type cells and *cap2*Δ mutant carrying GFP-Msb2p expressed from the *GAL* promoter. Cells were grown on semi-solid YEP-GAL media for 16 h. Scale bar, 10 microns. (**B**) Quantitation of cells in panel A. Error bars, standard error of mean from 3 independent trials. Asterisk, p-value < 0.05. (**C**) Colony immunoblot of indicated strains. Cells were spotted onto nitrocellulose membrane placed onto semi-solid YEP-GAL media and incubated at 30°C for 24 h. Colonies were washed off and the membrane was probed against anti-HA antibody.

The reticulated pattern of GFP-Msb2p localization seen in cells lacking Cap2p might reflect a problem with the trafficking of the protein. Msb2p is proteolytically processed in its N-terminal domain, and the processed extracellular glycodomain of the protein is secreted from the cell ^39, 40, 89^. We tested if the altered localization pattern of Msb2p in cells lacking Cap2p might impact Msb2p’s processing. An epitope-tagged version of Msb2p which contains the hemagglutinin (HA) epitope at amino acid 500 residues is shed from cells and was used to measure secretion of Msb2p by colony immunoblot analysis ^89^. The *cap2*Δ mutant showed elevated shedding of Msb2p, relative to wild-type cells and an fMAPK pathway mutant, *ste12*Δ (**Fig. 4C**). Thus, Msb2p was both trapped inside cells and shed at higher levels. One possibility is that in cells lacking Cap2p, Msb2p becomes trapped in intracellular compartments, which leads to increased proteolytic processing of the protein and elevated shedding of its extracellular domain. Indeed, Msb2p trapped in the secretory pathway is known to be processed efficiently ^39^.

Altered Msb2p trafficking in the *cap2*Δ mutant impacted the activity of the fMAPK pathway. The phosphorylation of the MAP kinase Kss1p (P∼Kss1p), the terminal kinase in the fMAPK cascade, occurs when the pathway is active. The level of P∼Kss1p provides a readout of fMAPK activity. P∼Kss1p levels showed a subtle signaling defect in the *cap2*Δ mutant compared to wild-type cells (**Fig. 5A**). Cells expressing a hyperactive version of Msb2p [*MSB2*^Δ100–818^]^40^ showed elevated fMAPK activity that was not reduced in cells lacking Cap2p (**Fig. 5A**). Expression of the *FUS1-lacZ* reporter was equivalent, based on statistical significance, between wild-type cells and the *cap2*Δ mutant (**Fig. 5B**). Growth of cells harboring the *FUS1-HIS3* reporter, which is a mating pathway reporter that in *ste4*Δ strains shows Msb2p-dependent fMAPK activity ^57^, showed a subtle signaling defect in the *cap2*Δ mutant (*Table 3*, fMAPK). Thus, Cap2p might play a minor role in regulating the activity of the fMAPK pathway. One possibility is that this occurs by impacting Msb2p’s trafficking. Nevertheless, this signaling defect is a subtle defect and we do not suggest that Cap2p is a component of the pathway.

**Figure 5.**
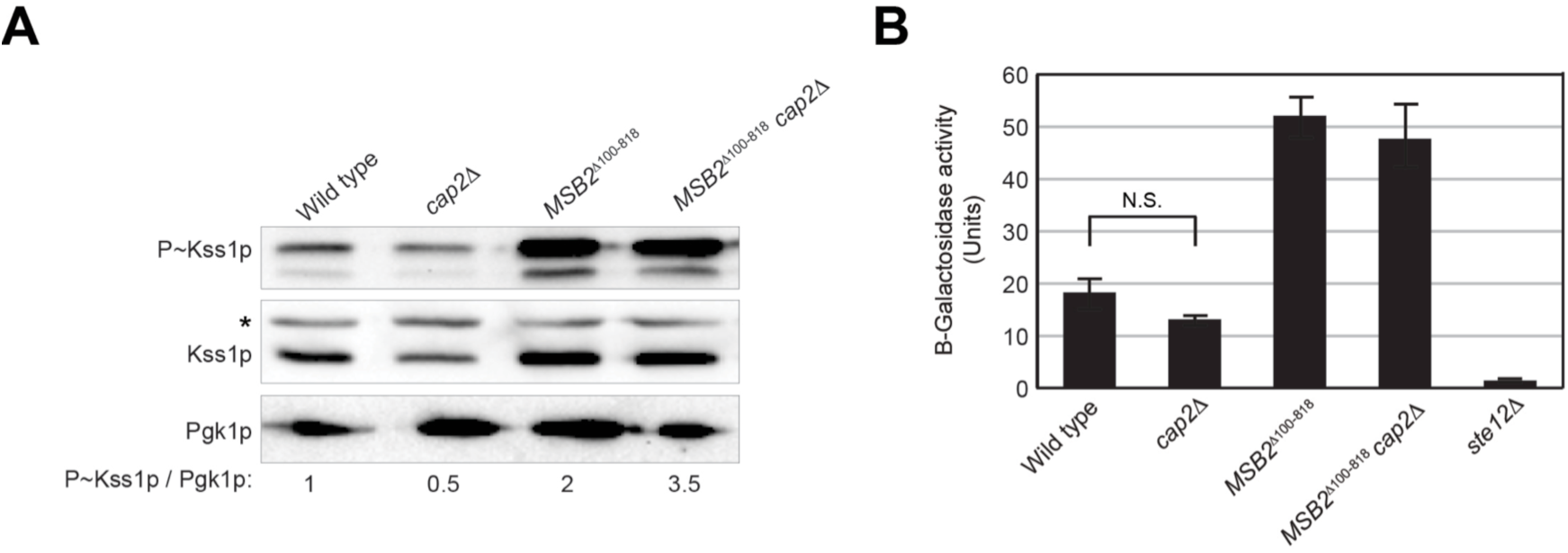
Role of Cap2p in regulating the fMAPK pathway. (**A**) The fMAPK pathway activity was compared among *cap2*Δ mutants in wild type and hyperactive Msb2p (*MSB2*^Δ100–818^) by measuring the levels of phosphorylated Kss1p. Asterisk, non-specific band. Numbers refer to ratio of P∼Kss1p to Pgk1p. (**B**) As for panel A, the activity of the fMAPK-dependent *FUS1-lacZ* reporter. The graph represents average of 3 trials. Error bars, standard error of mean. N.S., not significant.

Cap2p is an actin regulatory protein ^82, 90^. The actin cytoskeleton is highly polarized during filamentous growth to produce an elongated cell. Cap2p might impact filamentous growth through its role in regulating the actin cytoskeleton. The role that Cap2p played in regulating filamentous growth was assessed by the plate-washing assay ^30^ by measuring the agar invasion of the *cap2*Δ mutant relative to wild-type cells and control strains. The controls used were *msb2*Δ mutant and the *ste11*Δ mutant, which lacks the MAPKKK for the fMAPK pathway ^30^. Colonies were washed under a stream of water to reveal the invasion pattern as a read out of filamentous growth. The *cap2*Δ mutant was defective for invasive growth (**Fig. 6A**). The invasive growth defect of the *cap2*Δ mutant resembled the invasive growth defect of polarisome mutants (**Fig. 6A**, polarisome). These included cells lacking the formin Bni1p ^91^ or associated proteins Pea2p, Spa2p, and Bud6p in the polarisome ^92^, which has been previously shown to regulate invasive growth ^28, 93–95^. The polarisome also showed a minor role in fMAPK regulation as Cap2p (not shown). Given that the phenotype of the *cap2*Δ mutant in filamentous growth was similar to other polarity mutants, it might result from a general defect in cell polarity. Cells undergoing filamentous growth are highly polarized and more elongated than yeast-form cells ^26^. The *cap2*Δ mutant cells were rounder than wild-type cells under conditions that promote invasive growth (**Fig. 6B**, *cap2*Δ). Even in cells expressing hyperactive version of Msb2p that induces polarized growth, deletion of *CAP2* abolished the polarized morphology of cells (**Fig. 6B**, *MSB2*^Δ100–818^ *cap2*Δ).

**Figure 6.**
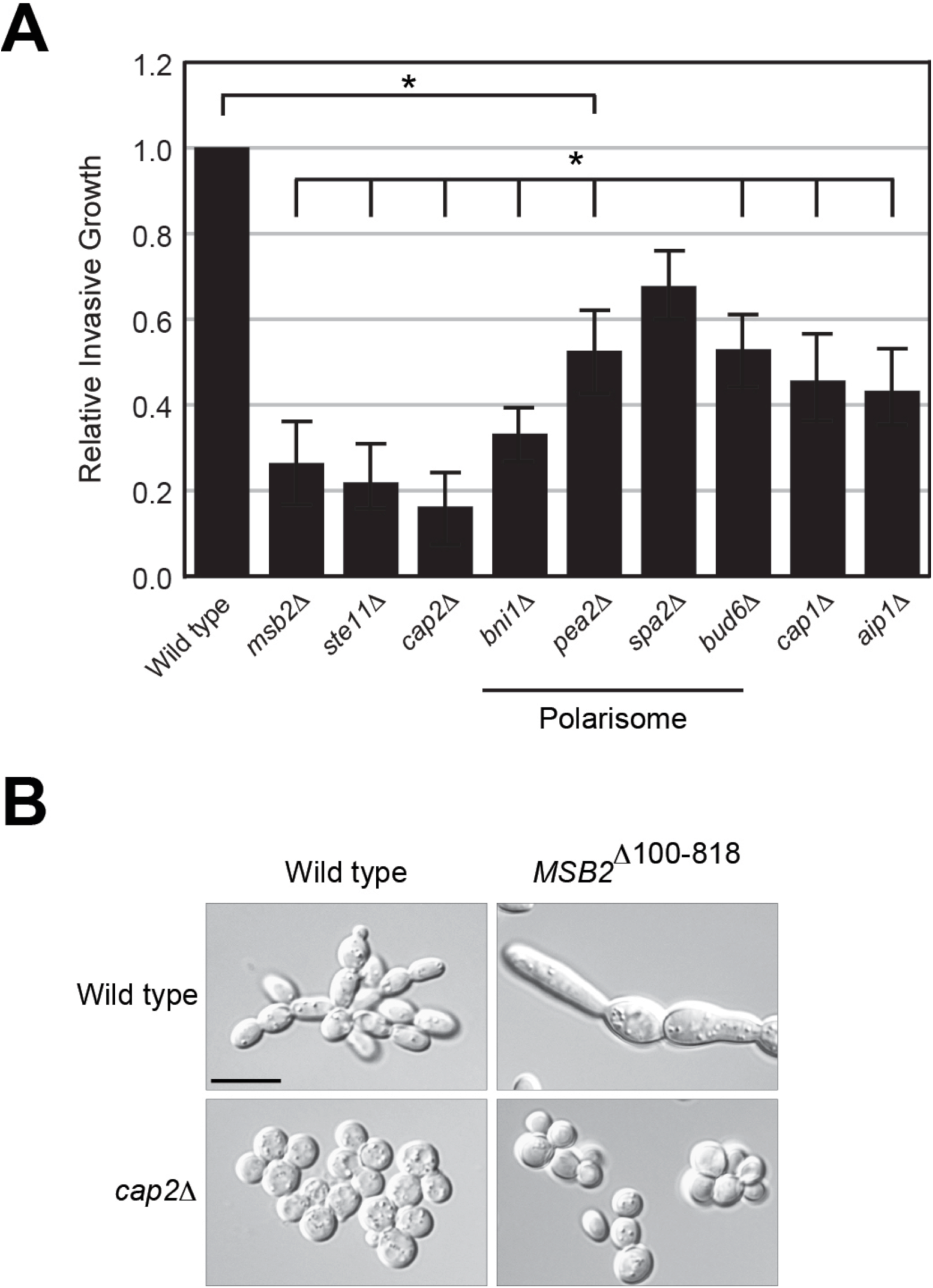
Cap2p regulates cell polarization during filamentous growth. (**A**) Invasive growth assay for indicated strains. Cells were spotted on YEPD medium. The plate was incubated at 30°C. At 3 d, the plates were photographed, washed in a stream of water, and photographed again to show invaded cells. Quantitation of invaded region was performed in ImageJ analysis by measuring the signal intensity for each strain normalized to wild type. Error bars, standard error of mean from two trials. Asterisk, p-value < 0.05. (**B**) The *cap2*Δ mutant is defective for the polarization of filamentous cells and for the polarization induced in cells containing an activated FG pathway. Wild-type cells, *cap2*Δ, *MSB2*^Δ100-818^, and *MSB2*^Δ100-818^ *cap2*Δ mutants were examined in YEP-GAL medium at 16 h. Representative cells are shown at 100X in DIC, bar, 10 microns.

Haploid cells undergoing filamentous growth switch to a distal-unipolar pattern ^26, 30, 96^. The *cap2*Δ mutant showed a defect in distal-pole budding under nutrient-limiting conditions (WT, Proximal: Equatorial: Distal = 34.13%: 64.42%: 1.45%; *cap2*Δ, Proximal: Equatorial: Distal = 75.59%: 19.25%: 5.16%). Under conditions in which cells bud axially, Cap2p did not impact budding pattern (WT, P:E:D 97.10%: 1.45%: 1.45%; *cap2*Δ, 95.12%: 2.93%: 1.95%). These results are consistent with the idea that when cells are less polarized they have less propensity for distal-pole budding ^97^ and may also account for the invasive growth defect of the *cap2*Δ mutant.

Cap2p heterodimerizes with Cap1p to form actin-capping protein. The *cap1*Δ mutant was phenotypically indistinguishable from the *cap2*Δ mutant by the PWA (**Fig. 6A**), which indicates that the actin capping function of Cap2p may account for its role in regulating cell polarization during filamentous growth. Like the *cap2*Δ mutant, the *cap1*Δ mutant showed a subtle defect in fMAPK signaling by *FUS1-HIS3* reporter (not shown).

Another actin-binding protein that interacted with Msb2p by protein microarray was Aip1p (*Table 2*). The *aip1*Δ mutant showed a similar phenotype as the *cap2*Δ mutant, in that it was defective for invasive growth by the PWA (**Fig. 4A**, *Table 3*). Thus, actin cytoskeletal regulatory proteins that interact with Msb2p might regulate filamentous growth by their roles in impacting Msb2p trafficking, and/or also by their established roles in regulating cell polarity.

Another Msb2p^CYT^ interacting protein, Nap1p, plays a role in chromatin organization and bud growth. Nap1p is a histone chaperone ^98, 99^ that also functions as a cell cycle regulatory protein. Nap1p is required for septin cytoskeleton organization ^100–102^ and a subset of mitotic events ^103, 104^ including regulation of the morphogenetic checkpoint ^105^. The plate-washing assay showed a role for Nap1p in invasive growth. Consistent with what has been shown in *Candida albicans* ^101^, the *S. cerevisiae nap1*Δ mutant was more invasive than wild-type cells (**Fig. 7A**). Examination of the cell morphology of the invaded cells showed a hyperpolarized growth phenotype. Nap1p did not regulate the fMAPK pathway, based on the activity of the *FUS1-HIS3* reporter (*Table 3*). Overexpression of *NAP1* induced polarized growth, concomitant with a growth defect, as has been previously reported ^106, 107^. The growth defect of *GAL-NAP1* cells was exacerbated by the loss of *MSB2* (**Fig. 7B**). Msb2p was also required for the hyperpolarized cell morphology of the *nap1*Δ mutant (**Fig. 7C**). These results demonstrate a genetic interaction between *MSB2* and *NAP1*.

**Figure 7.**
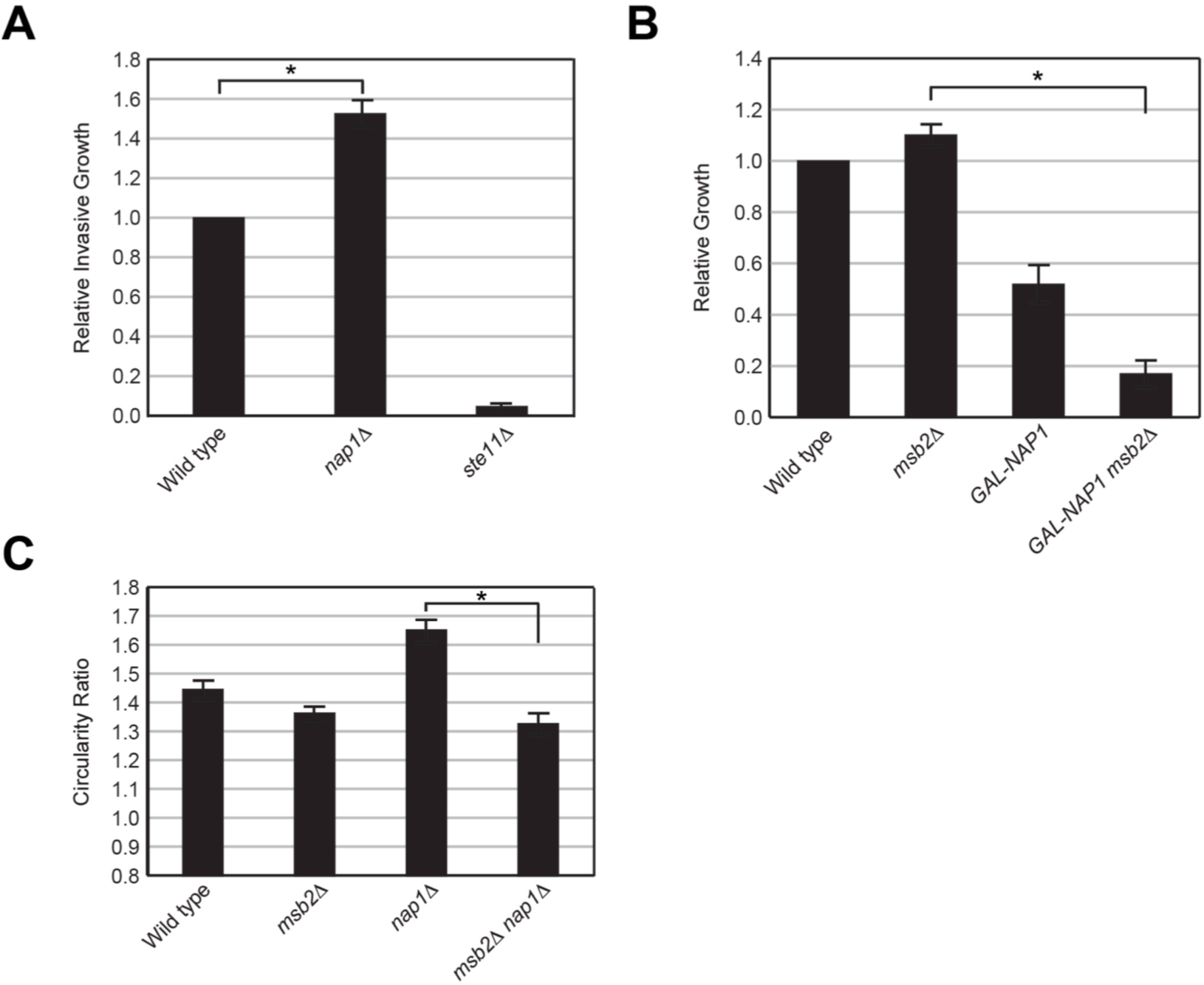
Functional analysis of *nap1*Δ mutant in filamentous growth. (**A**) The *nap1*Δ mutant was more invasive than wild-type cells by the PWA. See Fig 6A for details. (**B**) *MSB2* and *NAP1* show a genetic interaction. Serial dilutions of wild type, *msb2*Δ, p*GAL-NAP1* and p*GAL-NAP1 msb2*Δ cells were spotted onto S-GAL-URA plates. Equal concentrations were spotted on SD-URA as control. Growth of a colony was quantified by measuring the signal intensity of the colony against the background by ImageJ analysis. Growth on S-GAL-URA was compared to growth on SD-URA for each strain and then normalized to wild type. Error bars show the standard error of the mean from two separate trials. Asterisk, p-value < 0.05. (**C**) Polarized growth was evaluated for indicated strains. Cells were grown to saturation in YEPD media and visualized by DIC microscopy. For each cell, the circularity ratio was measured as the ratio of the long axis (distance from the mother bud neck to the distal pole) to the short axis (width of the cell) using ImageJ. Error bars, standard error of mean from two trials. 30 cells were measured in each trial. Asterisk, p-value < 0.05.

### Msb2p Shows Physical Interaction with Gln1p

The cytosolic domain of Msb2p showed interactions with other proteins (Table 2). One protein that we examined was Gln1p, which encodes glutamine synthetase ^108^. We validated this interaction by *in vitro* pull down analysis. Specifically, yeast cell extracts containing Gln1p-GFP were tested for interaction with beads bound to HIS-Msb2p^CYT^ purified in *E. coli*. Gln1p-GFP associated with HIS-Msb2p^CYT^ by *in vitro* pull down (**Fig. 8A**).

**Figure 8.**
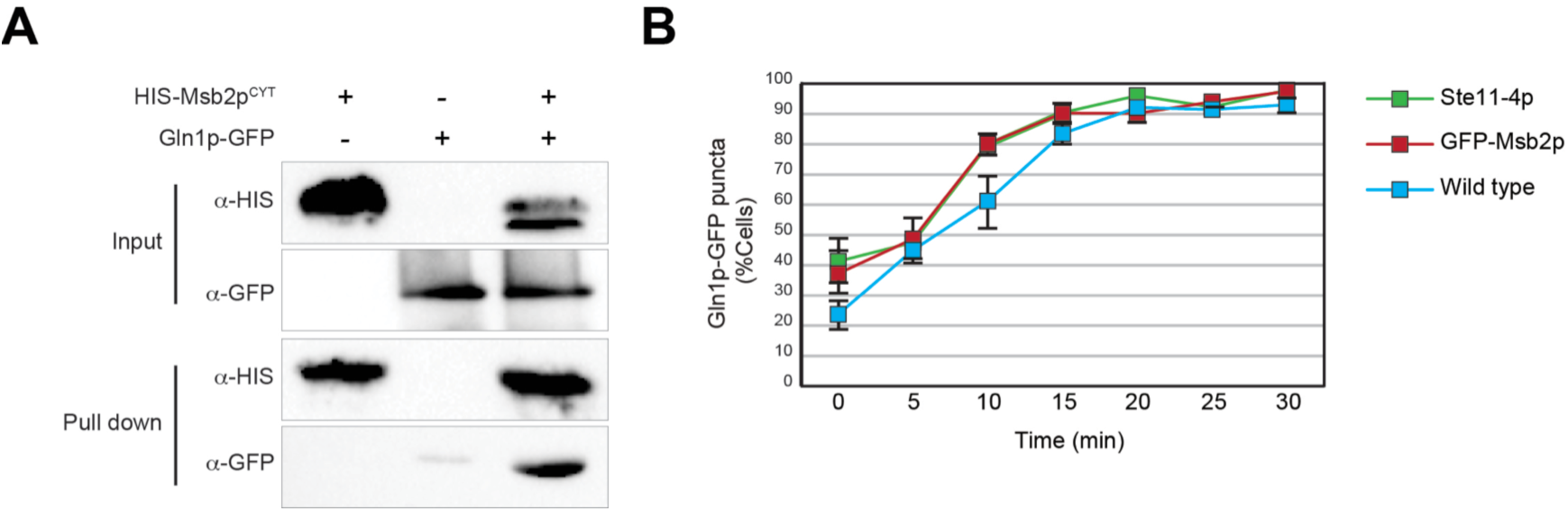
Msb2p interacts with Gln1p. (**A**) Msb2p interacts with glutamine synthetase, Gln1p, by *in vitro* pull down analysis. (**B**) The impact of Gln1p aggregation by the fMAPK pathway. Mid-log cells were induced by growth in citrate-phosphate buffer, pH 6.0 at t=0. Cells with visible GFP puncta were plotted as a percentage of total cells.

Glutamine is the key nitrogen precursor that feeds into several metabolic pathways that synthesize amino acids, purines and pyrimidines. As a result, the enzyme Gln1p that synthesizes glutamine is regulated at multiple levels inside the cell and its activity is sensitive to nutrient availability. Like other metabolic enzymes, one mode of regulation of Gln1p is its assembly into higher order inactive oligomers upon cellular starvation ^109, 110^. We tested if the interaction between Msb2p and Gln1p might impact Gln1p aggregation. Mid-log cells expressing Gln1p-GFP were switched to citrate-phosphate buffer, pH 6.0. Under this condition, Gln1p formed aggregates. Cells showing aggregation of Gln1p-GFP were similar among wild-type cells and cells expressing hyperactive alleles of Msb2p (GFP-Msb2p) or hyperactive alleles of Ste11p, the MAPKKK for the fMAPK pathway (**Fig. 8B**, Ste11-4p). Therefore, the interaction between Gln1p and Msb2p does not impact Gln1p aggregation.

## DISCUSSION

Here, protein microarrays were used to identify proteins that interact with the cytosolic domain of Msb2p, a mucin-like protein that regulates signal transduction (MAPK) pathways. One class of anticipated interacting partners were proteins that regulate MAPK pathways; however, such proteins were not identified by this approach. These interactions may be transient or require specific post-translational modifications on the cytosolic domain of Msb2p. The proteins identified by this approach have diverse biological functions. Some proteins might impact aspects of the life cycle of Msb2p. Others may connect Msb2p to diverse biological functions. For example, Msb2p may regulate the fMAPK pathway, which responds to nutrient availability, and directly modify the activity, localization, or function of metabolic enzymes. Metabolic enzymes can directly impact signaling pathway activity ^111^. Thus, the study provides a resource for interrogating the roles of the proteome in impacting mucin function and regulation in a model organism.

One connection that was explored in the study was between Msb2p and proteins that comprise the actin cytoskeleton. The actin cytoskeleton is a highly dynamic structure that controls diverse cellular functions ^112^. The fMAPK pathway is known to reorganize cell shape during filamentous growth, particularly by promoting distal-unipolar budding ^26, 96^ and causing a delay in the cell cycle that results in enhanced polarized growth by the polarisome ^27, 96^. Msb2p might directly regulate the actin cytoskeleton during filamentous growth through its interaction with actin binding proteins. In line with this possibility, two actin-binding proteins that interact with Msb2p, Cap2p and Aip1p, were required for the highly polarized cell morphologies that occurred during filamentous growth and for the subsequent invasive growth response. These proteins might be expected to regulate actin polymerization during filamentous growth, and whether Msb2p directly contributes to their function or activity in this context remains an open question. Msb2p through its role in the fMAPK pathway has been shown to regulate bud emergence and bud growth (Prabhakar et al submitted). Actin patches have also been shown to regulate polarized bud growth in the absence of actin cables ^113^. Thus, interactions between Msb2p and components of actin patches might reflect other roles for these proteins in regulation of polarized growth.

Actin capping protein is typically found associated with actin patches. In actin patches, Cap2p and other proteins play a role in endocytosis ^87^. Actin patches form a dynamic relationship with actin cables in yeast ^88^, and Cap2p and other proteins contribute to the transition between the different pools of actin in the cell ^90^. We show that Cap2p is required for the proper localization of Msb2p. One possibility is that Cap2p may impact the endocytosis of Msb2p from the plasma membrane. Msb2p is ubiquitinated at the plasma membrane and internalized by a mechanism that involves the ubiquitin ligase Rsp5p ^39^. In cells lacking Cap2p, Msb2p may reside longer at the plasma membrane, which might account for the elevated shedding of the protein. Msb2p shedding is regulated by yapsins, which proteolytically process Msb2p, and which are also known to function at the plasma membrane. It is also possible that Cap2p may impact the delivery of internalized Msb2p to other compartments in the secretory pathway, either to promote its recycling to the plasma membrane or its delivery to the lysosome. Like other mucins, Msb2p may be recycled to the plasma membrane, and the protein is known to be delivered by the ESCRT complex to the lysosome ^75^. In cells lacking Cap2p, Msb2p might become trapped in one or more of these compartments, resulting in elevated processing of the protein. This is supported by the mislocalization of Msb2p in cells lacking Cap2p. Yapsins are capable of processing Msb2p that has become trapped in the secretory pathway ^39^. Although processing of Msb2p is required for fMAPK pathway activation ^40^, the elevated processing of Msb2p did not stimulate Msb2p activity in this context, which suggests that this aspect of Msb2p regulation occurs after Msb2p-dpendent signaling has terminated. Intriguingly, mammalian mucins become progressively glycosylated through multiple rounds of delivery to the plasma membrane ^114^. There may be merit to exploring the roles of cytoskeletal proteins in regulating mucin trafficking in general. Actin filaments also act as barrier to MUC5AC secretion in lung cells, and disruption of the actin cytoskeleton by actin severing and capping proteins increases MUC5AC secretion ^115^. Thus, Msb2p shedding might be impacted by altered actin dynamics through an indirect mechanism from loss of Cap2p function.

We have also identified other protein interactions with Msb2p. One of these was with Nap1p, and another was with glutamine synthetase. Nitrogen levels are known to impact filamentous growth ^26^, including the levels of glutamine tRNA pools ^116, 117^. It is possible that Msb2p might impact the activity of glutamine synthetase to directly impact cellular metabolism. MUC1 expression increases glutamine levels in pancreatic cancer. ^118^. In addition, human MUC1 interacts with pyruvate kinase M2, which is the pyruvate kinase isoform found in cancer cells, to improve its activity ^119^. Future studies of the functionally relevant interactions between mucins and their interacting partners will likely provide new insights into functions of these important signaling regulatory proteins.

## ABBREVIATIONS

D: dextrose
Gal: galactose
GFP: green fluorescent protein
Glu: glucose
HA: hemaglutinin
HOG: high osmolarity glycerol response
co-IPT: co-immunoprecipitation; MAPK, mitogen activated protein kinase
PM: plasma membrane
SD: synthetic dextrose
WT: wild type

## ACKNOWLEDGEMENTS

Thanks to John Pringle (Stanford University), Charlie Boone (University of Toronto), Anuj Kumar (University of Michigan), and Mike Yu (University at Buffalo) for providing reagents. Thanks to Simon Alberti (Max Plank Institute of Molecular Cell Biology and Genetics, Dresden Germany) for helpful comments. The work was supported from a grant from the NIH (GM#098629). The authors declare no conflict of interests for the study.

## REFERENCES

[1] Bafna, S., Kaur, S., and Batra, S. K. (2010) Membrane-bound mucins: the mechanistic basis for alterations in the growth and survival of cancer cells, Oncogene 29, 2893–2904.

[2] Niv, Y., and Boltin, D. (2012) Secreted and membrane-bound mucins and idiopathic peptic ulcer disease, Digestion 86, 258–263.

[3] Birchenough, G. M., Johansson, M. E., Gustafsson, J. K., Bergstrom, J. H., and Hansson, G. C. (2015) New developments in goblet cell mucus secretion and function, Mucosal Immunol 8, 712–719.

[4] Singh, P. K., and Hollingsworth, M. A. (2006) Cell surface-associated mucins in signal transduction, Trends in cell biology 16, 467–476.

[5] Carraway, K. L., Ramsauer, V. P., Haq, B., and Carothers Carraway, C. A. (2003) Cell signaling through membrane mucins, BioEssays : news and reviews in molecular, cellular and developmental biology 25, 66–71.

[6] Kufe, D. W. (2009) Mucins in cancer: function, prognosis and therapy, Nat Rev Cancer 9, 874–885.

[7] Cross, B. W., and Ruhl, S. (2018) Glycan recognition at the saliva - oral microbiome interface, Cell Immunol 333, 19–33.

[8] Hollingsworth, M. A., and Swanson, B. J. (2004) Mucins in cancer: protection and control of the cell surface, Nature Reviews Cancer 4, 45–60.

[9] van Putten, J. P. M., and Strijbis, K. (2017) Transmembrane Mucins: Signaling Receptors at the Intersection of Inflammation and Cancer, J Innate Immun 9, 281–299.

[10] Xu, M., Wang, D. C., Wang, X., and Zhang, Y. (2017) Correlation between mucin biology and tumor heterogeneity in lung cancer, Seminars in cell & developmental biology 64, 73–78.

[11] Kufe, D. W. (2013) MUC1-C oncoprotein as a target in breast cancer: activation of signaling pathways and therapeutic approaches, Oncogene 32, 1073–1081.

[12] Moniaux, N., Escande, F., Porchet, N., Aubert, J. P., and Batra, S. K. (2001) Structural organization and classification of the human mucin genes, Front Biosci 6, D1192–1206.

[13] Chaturvedi, P., Singh, A. P., and Batra, S. K. (2008) Structure, evolution, and biology of the MUC4 mucin, FASEB journal : official publication of the Federation of American Societies for Experimental Biology 22, 966–981.

[14] Gendler, S. J., Burchell, J. M., Duhig, T., Lamport, D., White, R., Parker, M., and Taylor-Papadimitriou, J. (1987) Cloning of partial cDNA encoding differentiation and tumor-associated mucin glycoproteins expressed by human mammary epithelium, Proc Natl Acad Sci U S A 84, 6060–6064.

[15] Timpte, C. S., Eckhardt, A. E., Abernethy, J. L., and Hill, R. L. (1988) Porcine submaxillary gland apomucin contains tandemly repeated, identical sequences of 81 residues, J Biol Chem 263, 1081–1088.

[16] Gupta, R., and Jentoft, N. (1989) Subunit structure of porcine submaxillary mucin, Biochemistry 28, 6114–6121.

[17] Silverman, H. S., Sutton-Smith, M., McDermott, K., Heal, P., Leir, S. H., Morris, H. R., Hollingsworth, M. A., Dell, A., and Harris, A. (2003) The contribution of tandem repeat number to the O-glycosylation of mucins, Glycobiology 13, 265–277.

[18] Tian, E., and Ten Hagen, K. G. (2009) Recent insights into the biological roles of mucin-type O-glycosylation, Glycoconj J 26, 325–334.

[19] Tarp, M. A., and Clausen, H. (2008) Mucin-type O-glycosylation and its potential use in drug and vaccine development, Biochimica et biophysica acta 1780, 546–563.

[20] Cullen, P. J., Sabbagh, W., Jr., Graham, E., Irick, M. M., van Olden, E. K., Neal, C., Delrow, J., Bardwell, L., and Sprague, G. F., Jr. (2004) A signaling mucin at the head of the Cdc42- and MAPK-dependent filamentous growth pathway in yeast, Genes & development 18, 1695–1708.

[21] Clevers, H. (2004) Signaling mucins in the (S)limelight, Developmental cell 7, 150–151.

[22] Gancedo, J. M. (2001) Control of pseudohyphae formation in Saccharomyces cerevisiae, FEMS Microbiol Rev 25, 107–123.

[23] Pan, X., Harashima, T., and Heitman, J. (2000) Signal transduction cascades regulating pseudohyphal differentiation of Saccharomyces cerevisiae, Current opinion in microbiology 3, 567–572.

[24] Wendland, J. (2001) Comparison of morphogenetic networks of filamentous fungi and yeast, Fungal genetics and biology : FG & B 34, 63–82.

[25] Whiteway, M., and Bachewich, C. (2007) Morphogenesis in Candida albicans, Annual review of microbiology 61, 529–553.

[26] Gimeno, C. J., Ljungdahl, P. O., Styles, C. A., and Fink, G. R. (1992) Unipolar cell divisions in the yeast S. cerevisiae lead to filamentous growth: regulation by starvation and RAS, Cell 68, 1077–1090.

[27] Kron, S. J., Styles, C. A., and Fink, G. R. (1994) Symmetric cell division in pseudohyphae of the yeast Saccharomyces cerevisiae, Molecular biology of the cell 5, 1003–1022.

[28] Rua, D., Tobe, B. T., and Kron, S. J. (2001) Cell cycle control of yeast filamentous growth, Current opinion in microbiology 4, 720–727.

[29] Verstrepen, K. J., and Klis, F. M. (2006) Flocculation, adhesion and biofilm formation in yeasts, Molecular microbiology 60, 5–15.

[30] Roberts, R. L., and Fink, G. R. (1994) Elements of a single MAP kinase cascade in Saccharomyces cerevisiae mediate two developmental programs in the same cell type: mating and invasive growth, Genes & development 8, 2974–2985.

[31] Borneman, A. R., Gianoulis, T. A., Zhang, Z. D., Yu, H., Rozowsky, J., Seringhaus, M. R., Wang, L. Y., Gerstein, M., and Snyder, M. (2007) Divergence of transcription factor binding sites across related yeast species, Science 317, 815–819.

[32] Leberer, E., Wu, C., Leeuw, T., Fourest-Lieuvin, A., Segall, J. E., and Thomas, D. Y. (1997) Functional characterization of the Cdc42p binding domain of yeast Ste20p protein kinase, The EMBO journal 16, 83–97.

[33] Peter, M., Neiman, A. M., Park, H. O., van Lohuizen, M., and Herskowitz, I. (1996) Functional analysis of the interaction between the small GTP binding protein Cdc42 and the Ste20 protein kinase in yeast, The EMBO journal 15, 7046–7059.

[34] Madhani, H. D., Styles, C. A., and Fink, G. R. (1997) MAP kinases with distinct inhibitory functions impart signaling specificity during yeast differentiation, Cell 91, 673–684.

[35] Madhani, H. D., and Fink, G. R. (1997) Combinatorial control required for the specificity of yeast MAPK signaling, Science 275, 1314–1317.

[36] Rupp, S., Summers, E., Lo, H. J., Madhani, H., and Fink, G. (1999) MAP kinase and cAMP filamentation signaling pathways converge on the unusually large promoter of the yeast FLO11 gene, The EMBO journal 18, 1257–1269.

[37] Roberts, C. J., Nelson, B., Marton, M. J., Stoughton, R., Meyer, M. R., Bennett, H. A., He, Y. D., Dai, H., Walker, W. L., Hughes, T. R., Tyers, M., Boone, C., and Friend, S. H. (2000) Signaling and circuitry of multiple MAPK pathways revealed by a matrix of global gene expression profiles, Science 287, 873–880.

[38] Lien, E. C., Nagiec, M. J., and Dohlman, H. G. (2013) Proper protein glycosylation promotes mitogen-activated protein kinase signal fidelity, Biochemistry 52, 115–124.

[39] Adhikari, H., Vadaie, N., Chow, J., Caccamise, L. M., Chavel, C. A., Li, B., Bowitch, A., Stefan, C. J., and Cullen, P. J. (2015) Role of the unfolded protein response in regulating the mucin-dependent filamentous-growth mitogen-activated protein kinase pathway, Molecular and cellular biology 35, 1414–1432.

[40] Vadaie, N., Dionne, H., Akajagbor, D. S., Nickerson, S. R., Krysan, D. J., and Cullen, P. J. (2008) Cleavage of the signaling mucin Msb2 by the aspartyl protease Yps1 is required for MAPK activation in yeast, The Journal of cell biology 181, 1073–1081.

[41] Tatebayashi, K., Tanaka, K., Yang, H. Y., Yamamoto, K., Matsushita, Y., Tomida, T., Imai, M., and Saito, H. (2007) Transmembrane mucins Hkr1 and Msb2 are putative osmosensors in the SHO1 branch of yeast HOG pathway, The EMBO journal 26, 3521–3533.

[42] Pitoniak, A., Birkaya, B., Dionne, H. M., Vadaie, N., and Cullen, P. J. (2009) The signaling mucins Msb2 and Hkr1 differentially regulate the filamentation mitogen-activated protein kinase pathway and contribute to a multimodal response, Molecular biology of the cell 20, 3101–3114.

[43] O’Rourke, S. M., and Herskowitz, I. (2002) A third osmosensing branch in Saccharomyces cerevisiae requires the Msb2 protein and functions in parallel with the Sho1 branch, Molecular and cellular biology 22, 4739–4749.

[44] Tanaka, K., Tatebayashi, K., Nishimura, A., Yamamoto, K., Yang, H. Y., and Saito, H. (2014) Yeast osmosensors Hkr1 and Msb2 activate the Hog1 MAPK cascade by different mechanisms, Science signaling 7, ra21.

[45] Gelperin, D. M., White, M. A., Wilkinson, M. L., Kon, Y., Kung, L. A., Wise, K. J., Lopez-Hoyo, N., Jiang, L., Piccirillo, S., Yu, H., Gerstein, M., Dumont, M. E., Phizicky, E. M., Snyder, M., and Grayhack, E. J. (2005) Biochemical and genetic analysis of the yeast proteome with a movable ORF collection, Genes & development 19, 2816–2826.

[46] Baudin, A., Ozier-Kalogeropoulos, O., Denouel, A., Lacroute, F., and Cullin, C. (1993) A simple and efficient method for direct gene deletion in Saccharomyces cerevisiae, Nucleic Acids Res 21, 3329–3330.

[47] Longtine, M. S., McKenzie, A., 3rd, Demarini, D. J., Shah, N. G., Wach, A., Brachat, A., Philippsen, P., and Pringle, J. R. (1998) Additional modules for versatile and economical PCR-based gene deletion and modification in Saccharomyces cerevisiae, Yeast 14, 953–961.

[48] Goldstein, A. L., and McCusker, J. H. (1999) Three new dominant drug resistance cassettes for gene disruption in Saccharomyces cerevisiae, Yeast 15, 1541–1553.

[49] Schneider, B. L., Seufert, W., Steiner, B., Yang, Q. H., and Futcher, A. B. (1995) Use of polymerase chain reaction epitope tagging for protein tagging in Saccharomyces cerevisiae, Yeast 11, 1265–1274.

[50] Ghaemmaghami, S., Huh, W. K., Bower, K., Howson, R. W., Belle, A., Dephoure, N., O’Shea, E. K., and Weissman, J. S. (2003) Global analysis of protein expression in yeast, Nature 425, 737–741.

[51] Huh, W. K., Falvo, J. V., Gerke, L. C., Carroll, A. S., Howson, R. W., Weissman, J. S., and O’Shea, E. K. (2003) Global analysis of protein localization in budding yeast, Nature 425, 686–691.

[52] Sikorski, R. S., and Hieter, P. (1989) A system of shuttle vectors and yeast host strains designed for efficient manipulation of DNA in Saccharomyces cerevisiae, Genetics 122, 19–27.

[53] Stevenson, B. J., Rhodes, N., Errede, B., and Sprague, G. F., Jr. (1992) Constitutive mutants of the protein kinase STE11 activate the yeast pheromone response pathway in the absence of the G protein, Genes & development 6, 1293–1304.

[54] Sambrook, J., Fritsch, E.F., and Maniatis, T. (1989) Molecular cloning: a laboratory manual., Cold Spring Harbor Laboratory Press, Cold Spring Harbor, NY.

[55] Rose, M. D., Winston, F., and Hieter, P. (1990) Methods in yeast genetics., Cold Spring Harbor Laboratory Press, Cold Spring Harbor, NY.

[56] Cullen, P. J., Schultz, J., Horecka, J., Stevenson, B. J., Jigami, Y., and Sprague, G. F., Jr. (2000) Defects in protein glycosylation cause SHO1-dependent activation of a STE12 signaling pathway in yeast, Genetics 155, 1005–1018.

[57] McCaffrey, G., Clay, F. J., Kelsay, K., and Sprague, G. F., Jr. (1987) Identification and regulation of a gene required for cell fusion during mating of the yeast Saccharomyces cerevisiae, Molecular and cellular biology 7, 2680–2690.

[58] Cullen, P. J., and Sprague, G. F., Jr. (2000) Glucose depletion causes haploid invasive growth in yeast, Proceedings of the National Academy of Sciences of the United States of America 97, 13619–13624.

[59] Chant, J., and Pringle, J. R. (1995) Patterns of bud-site selection in the yeast Saccharomyces cerevisiae, The Journal of cell biology 129, 751–765.

[60] Sabbagh, W., Jr., Flatauer, L. J., Bardwell, A. J., and Bardwell, L. (2001) Specificity of MAP kinase signaling in yeast differentiation involves transient versus sustained MAPK activation, Molecular cell 8, 683–691.

[61] Lee, M. J., and Dohlman, H. G. (2008) Coactivation of G protein signaling by cell-surface receptors and an intracellular exchange factor, Current biology : CB 18, 211–215.

[62] Basu, S., Vadaie, N., Prabhakar, A., Li, B., Adhikari, H., Pitoniak, A., Chow, J., Chavel, C. A., and Cullen, P. J. (2016) Spatial landmarks regulate a Cdc42-dependent MAPK pathway to control differentiation and the response to positional compromise, Proceedings of the National Academy of Sciences of the United States of America 113, E2019–2028.

[63] Chavel, C. A., Dionne, H. M., Birkaya, B., Joshi, J., and Cullen, P. J. (2010) Multiple signals converge on a differentiation MAPK pathway, PLoS genetics 6, e1000883.

[64] Zhu, H., Klemic, J. F., Chang, S., Bertone, P., Casamayor, A., Klemic, K. G., Smith, D., Gerstein, M., Reed, M. A., and Snyder, M. (2000) Analysis of yeast protein kinases using protein chips, Nat Genet 26, 283–289.

[65] Kemp, H. A., and Sprague, G. F., Jr. (2003) Far3 and five interacting proteins prevent premature recovery from pheromone arrest in the budding yeast Saccharomyces cerevisiae, Molecular and cellular biology 23, 1750–1763.

[66] Zhu, H., Bilgin, M., Bangham, R., Hall, D., Casamayor, A., Bertone, P., Lan, N., Jansen, R., Bidlingmaier, S., Houfek, T., Mitchell, T., Miller, P., Dean, R. A., Gerstein, M., and Snyder, M. (2001) Global analysis of protein activities using proteome chips, Science 293, 2101–2105.

[67] Zhu, Z. M., and Wang, X. Q. (1998) Role for cell surface oligosaccharide in cell-cell recognition during implantation, Mol Hum Reprod 4, 735–738.

[68] Neiswinger, J., Uzoma, I., Cox, E., Rho, H., Song, G., Paul, C., Jeong, J. S., Lu, K. Y., Chen, C. S., and Zhu, H. (2016) Protein Microarrays: Flexible Tools for Scientific Innovation, Cold Spring Harbor protocols 2016.

[69] Paul, C., Rho, H., Neiswinger, J., and Zhu, H. (2016) Characterization of Protein-Protein Interactions Using Protein Microarrays, Cold Spring Harbor protocols 2016.

[70] Lu, J. Y., Lin, Y. Y., Boeke, J. D., and Zhu, H. (2013) Using functional proteome microarrays to study protein lysine acetylation, Methods in molecular biology 981, 151–165.

[71] Tao, S. C., Li, Y., Zhou, J., Qian, J., Schnaar, R. L., Zhang, Y., Goldstein, I. J., Zhu, H., and Schneck, J. P. (2008) Lectin microarrays identify cell-specific and functionally significant cell surface glycan markers, Glycobiology 18, 761–769.

[72] Hu, S., Xie, Z., Blackshaw, S., Qian, J., and Zhu, H. (2011) Characterization of protein-DNA interactions using protein microarrays, Cold Spring Harbor protocols 2011, pdb prot5614.

[73] Song, G., Neiswinger, J., and Zhu, H. (2016) Characterization of RNA-Binding Proteins Using Protein Microarrays, Cold Spring Harbor protocols 2016.

[74] Herianto, S., Chen, C. S., and Zhu, H. (2019) Protein Microarrays and Liposome: A Method for Studying Lipid-Protein Interactions, Methods in molecular biology 2003, 191–199.

[75] Adhikari, H., Caccamise, L. M., Pande, T., and Cullen, P. J. (2015) Comparative Analysis of Transmembrane Regulators of the Filamentous Growth Mitogen-Activated Protein Kinase Pathway Uncovers Functional and Regulatory Differences, Eukaryotic cell 14, 868–883.

[76] Karunanithi, S., and Cullen, P. J. (2012) The Filamentous Growth MAPK Pathway Responds to Glucose Starvation Through the Mig1/2 Transcriptional Repressors in Saccharomyces cerevisiae, Genetics 192, 869–887.

[77] Eden, E., Navon, R., Steinfeld, I., Lipson, D., and Yakhini, Z. (2009) GOrilla: a tool for discovery and visualization of enriched GO terms in ranked gene lists, BMC Bioinformatics 10, 48.

[78] Eden, E., Lipson, D., Yogev, S., and Yakhini, Z. (2007) Discovering motifs in ranked lists of DNA sequences, PLoS Comput Biol 3, e39.

[79] Ryan, O., Shapiro, R. S., Kurat, C. F., Mayhew, D., Baryshnikova, A., Chin, B., Lin, Z. Y., Cox, M. J., Vizeacoumar, F., Cheung, D., Bahr, S., Tsui, K., Tebbji, F., Sellam, A., Istel, F., Schwarzmuller, T., Reynolds, T. B., Kuchler, K., Gifford, D. K., Whiteway, M., Giaever, G., Nislow, C., Costanzo, M., Gingras, A. C., Mitra, R. D., Andrews, B., Fink, G. R., Cowen, L. E., and Boone, C. (2012) Global gene deletion analysis exploring yeast filamentous growth, Science 337, 1353–1356.

[80] Yang, H. Y., Tatebayashi, K., Yamamoto, K., and Saito, H. (2009) Glycosylation defects activate filamentous growth Kss1 MAPK and inhibit osmoregulatory Hog1 MAPK, The EMBO journal 28, 1380–1391.

[81] Yamamoto, K., Tatebayashi, K., and Saito, H. (2015) Binding of the Extracellular Eight-Cysteine Motif of Opy2 to the Putative Osmosensor Msb2 Is Essential for Activation of the Yeast High-Osmolarity Glycerol Pathway, Molecular and cellular biology 36, 475–487.

[82] Amatruda, J. F., and Cooper, J. A. (1992) Purification, characterization, and immunofluorescence localization of Saccharomyces cerevisiae capping protein, The Journal of cell biology 117, 1067–1076.

[83] Amatruda, J. F., Cannon, J. F., Tatchell, K., Hug, C., and Cooper, J. A. (1990) Disruption of the actin cytoskeleton in yeast capping protein mutants, Nature 344, 352–354.

[84] Muhua, L., Karpova, T. S., and Cooper, J. A. (1994) A yeast actin-related protein homologous to that in vertebrate dynactin complex is important for spindle orientation and nuclear migration, Cell 78, 669–679.

[85] Bruck, S., Huber, T. B., Ingham, R. J., Kim, K., Niederstrasser, H., Allen, P. M., Pawson, T., Cooper, J. A., and Shaw, A. S. (2006) Identification of a novel inhibitory actin-capping protein binding motif in CD2-associated protein, The Journal of biological chemistry 281, 19196–19203.

[86] Hernandez-Valladares, M., Kim, T., Kannan, B., Tung, A., Aguda, A. H., Larsson, M., Cooper, J. A., and Robinson, R. C. (2010) Structural characterization of a capping protein interaction motif defines a family of actin filament regulators, Nat Struct Mol Biol 17, 497–503.

[87] Kaksonen, M., Toret, C. P., and Drubin, D. G. (2005) A modular design for the clathrin- and actin-mediated endocytosis machinery, Cell 123, 305–320.

[88] Berro, J., and Pollard, T. D. (2014) Synergies between Aip1p and capping protein subunits (Acp1p and Acp2p) in clathrin-mediated endocytosis and cell polarization in fission yeast, Molecular biology of the cell 25, 3515–3527.

[89] Chavel, C. A., Dionne, H. M., Birkaya, B., Joshi, J., and Cullen, P. J. (2010) Multiple Signals Converge on a Differentiation MAPK Pathway, PLOS Genetics 6, e1000883.

[90] Shin, M., van Leeuwen, J., Boone, C., and Bretscher, A. (2018) Yeast Aim21/Tda2 both regulates free actin by reducing barbed end assembly and forms a complex with Cap1/Cap2 to balance actin assembly between patches and cables, Molecular biology of the cell 29, 923–936.

[91] Evangelista, M., Blundell, K., Longtine, M. S., Chow, C. J., Adames, N., Pringle, J. R., Peter, M., and Boone, C. (1997) Bni1p, a yeast formin linking cdc42p and the actin cytoskeleton during polarized morphogenesis, Science 276, 118–122.

[92] Bidlingmaier, S., and Snyder, M. (2004) Regulation of polarized growth initiation and termination cycles by the polarisome and Cdc42 regulators, The Journal of cell biology 164, 207–218.

[93] Mosch, H. U., and Fink, G. R. (1997) Dissection of filamentous growth by transposon mutagenesis in Saccharomyces cerevisiae, Genetics 145, 671–684.

[94] Kang, C. M., and Jiang, Y. W. (2005) Genome-wide survey of non-essential genes required for slowed DNA synthesis-induced filamentous growth in yeast, *Yeast (Chichester*, England) 22, 79–90.

[95] Jin, R., Dobry, C. J., McCown, P. J., and Kumar, A. (2008) Large-scale analysis of yeast filamentous growth by systematic gene disruption and overexpression, Molecular biology of the cell 19, 284–296.

[96] Cullen, P. J., and Sprague, G. F., Jr. (2002) The roles of bud-site-selection proteins during haploid invasive growth in yeast, Molecular biology of the cell 13, 2990–3004.

[97] Sheu, Y. J., Barral, Y., and Snyder, M. (2000) Polarized growth controls cell shape and bipolar bud site selection in Saccharomyces cerevisiae, Molecular and cellular biology 20, 5235–5247.

[98] Ishimi, Y., and Kikuchi, A. (1991) Identification and molecular cloning of yeast homolog of nucleosome assembly protein I which facilitates nucleosome assembly in vitro, J Biol Chem 266, 7025–7029.

[99] Mosammaparast, N., Ewart, C. S., and Pemberton, L. F. (2002) A role for nucleosome assembly protein 1 in the nuclear transport of histones H2A and H2B, Embo j 21, 6527–6538.

[100] Longtine, M. S., Theesfeld, C. L., McMillan, J. N., Weaver, E., Pringle, J. R., and Lew, D. J. (2000) Septin-dependent assembly of a cell cycle-regulatory module in Saccharomyces cerevisiae, Molecular and cellular biology 20, 4049–4061.

[101] Huang, Z.-X., Zhao, P., Zeng, G.-S., Wang, Y.-M., Sudbery, I., and Wang, Y. (2014) Phosphoregulation of Nap1 Plays a Role in Septin Ring Dynamics and Morphogenesis in Candida albicans, mBio 5.

[102] Gladfelter, A. S., Zyla, T. R., and Lew, D. J. (2004) Genetic Interactions among Regulators of Septin Organization, Eukaryotic Cell 3, 847–854.

[103] Kellogg, D. R., and A.W. Murray. (1995) NAP1 acts with Clb1 to perform mitotic functions and to suppress polar bud growth in budding yeast, The Journal of Cell Biology 130, 675–685.

[104] Altman, R., and Kellogg, D. (1997) Control of Mitotic Events by Nap1 and the Gin4 Kinase, The Journal of Cell Biology 138, 119–130.

[105] Lew, D. J., and Reed, S. I. (1995) A cell cycle checkpoint monitors cell morphogenesis in budding yeast, The Journal of cell biology 129, 739–749.

[106] Calvert, M. E., Lannigan, J. A., and Pemberton, L. F. (2008) Optimization of yeast cell cycle analysis and morphological characterization by multispectral imaging flow cytometry, *Cytometry.* Part A : the journal of the International Society for Analytical Cytology 73, 825–833.

[107] Yoshikawa, K., Tanaka, T., Ida, Y., Furusawa, C., Hirasawa, T., and Shimizu, H. (2011) Comprehensive phenotypic analysis of single-gene deletion and overexpression strains of Saccharomyces cerevisiae, *Yeast (Chichester*, England) 28, 349–361.

[108] Mitchell, A. P., and Magasanik, B. (1984) Biochemical and physiological aspects of glutamine synthetase inactivation in Saccharomyces cerevisiae, The Journal of biological chemistry 259, 12054–12062.

[109] Narayanaswamy, R., Levy, M., Tsechansky, M., Stovall, G. M., O’Connell, J. D., Mirrielees, J., Ellington, A. D., and Marcotte, E. M. (2009) Widespread reorganization of metabolic enzymes into reversible assemblies upon nutrient starvation, Proceedings of the National Academy of Sciences 106, 10147–10152.

[110] Petrovska, I., Nuske, E., Munder, M. C., Kulasegaran, G., Malinovska, L., Kroschwald, S., Richter, D., Fahmy, K., Gibson, K., Verbavatz, J. M., and Alberti, S. (2014) Filament formation by metabolic enzymes is a specific adaptation to an advanced state of cellular starvation, eLife.

[111] Hall, D. A., Zhu, H., Zhu, X., Royce, T., Gerstein, M., and Snyder, M. (2004) Regulation of gene expression by a metabolic enzyme, Science 306, 482–484.

[112] Haarer, B., Viggiano, S., Hibbs, M. A., Troyanskaya, O. G., and Amberg, D. C. (2007) Modeling complex genetic interactions in a simple eukaryotic genome: actin displays a rich spectrum of complex haploinsufficiencies, Genes & development 21, 148–159.

[113] Yamamoto, T., Mochida, J., Kadota, J., Takeda, M., Bi, E., and Tanaka, K. (2010) Initial polarized bud growth by endocytic recycling in the absence of actin cable-dependent vesicle transport in yeast, Mol Biol Cell 21, 1237–1252.

[114] Engelmann, K., Kinlough, C. L., Muller, S., Razawi, H., Baldus, S. E., Hughey, R. P., and Hanisch, F. G. (2005) Transmembrane and secreted MUC1 probes show trafficking-dependent changes in O-glycan core profiles, Glycobiology 15, 1111–1124.

[115] Ehre, C., Rossi, A. H., Abdullah, L. H., De Pestel, K., Hill, S., Olsen, J. C., and Davis, C. W. (2005) Barrier role of actin filaments in regulated mucin secretion from airway goblet cells, American journal of physiology. Cell physiology 288, C46–56.

[116] Murray, L. E., Rowley, N., Dawes, I. W., Johnston, G. C., and Singer, R. A. (1998) A yeast glutamine tRNA signals nitrogen status for regulation of dimorphic growth and sporulation, Proceedings of the National Academy of Sciences of the United States of America 95, 8619–8624.

[117] Bjork, G. R., Huang, B., Persson, O. P., and Bystrom, A. S. (2007) A conserved modified wobble nucleoside (mcm5s2U) in lysyl-tRNA is required for viability in yeast, *RNA (New York*, N.Y.) 13, 1245–1255.

[118] Chaika, N. V., Gebregiworgis, T., Lewallen, M. E., Purohit, V., Radhakrishnan, P., Liu, X., Zhang, B., Mehla, K., Brown, R. B., Caffrey, T., Yu, F., Johnson, K. R., Powers, R., Hollingsworth, M. A., and Singh, P. K. (2012) MUC1 mucin stabilizes and activates hypoxia-inducible factor 1 alpha to regulate metabolism in pancreatic cancer, Proceedings of the National Academy of Sciences of the United States of America 109, 13787–13792.

[119] Kosugi, M., Ahmad, R., Alam, M., Uchida, Y., and Kufe, D. (2011) MUC1-C oncoprotein regulates glycolysis and pyruvate kinase M2 activity in cancer cells, PloS one 6, e28234.

## REFERENCES

[1] Cullen, P. J., and Sprague, G. F., Jr. (2002) The roles of bud-­-site-­-selection proteins during haploid invasive growth in yeast, Mol Biol Cell 13, 2990–3004.

[2] Cullen, P. J., Sabbagh, W., Jr., Graham, E., Irick, M. M., van Olden, E. K., Neal, C., Delrow, J., Bardwell, L., and Sprague, G. F., Jr. (2004) A signaling mucin at the head of the Cdc42-­- and MAPK-­-dependent filamentous growth pathway in yeast, Genes Dev 18, 1695–1708.

[3] Vadaie, N., Dionne, H., Akajagbor, D. S., Nickerson, S. R., Krysan, D. J., and Cullen, P. J. (2008) Cleavage of the signaling mucin Msb2 by the aspartyl protease Yps1 is required for MAPK activation in yeast, J Cell Biol 181, 1073–1081.

[4] Ghaemmaghami, S., Huh, W. K., Bower, K., Howson, R. W., Belle, A., Dephoure, N., O’Shea, E. K., and Weissman, J. S. (2003) Global analysis of protein expression in yeast, Nature 425, 737–741.

[5] Huh, W. K., Falvo, J. V., Gerke, L. C., Carroll, A. S., Howson, R. W., Weissman, J. S., and O’Shea, E. K. (2003) Global analysis of protein localization in budding yeast, Nature 425, 686–691.

